# Repurposing cell growth-regulating compounds identifies kenpaullone which ameliorates pathologic pain via normalization of inhibitory neurotransmission

**DOI:** 10.1101/767681

**Authors:** Michele Yeo, Yong Chen, Changyu Jiang, Gang Chen, Kaiyuan Wang, Andrey Bortsov, Maria Lioudyno, Qian Zeng, Zilong Wang, Jorge Busciglio, Ru-Rong Ji, Wolfgang Liedtke

**Author notes:** lead contact & corresponding author.

## Abstract

Inhibitory GABA-ergic neurotransmission is fundamental for the adult vertebrate central nervous system and requires low chloride ion concentration in neurons. This basic ionic-homeostatic mechanism relies on expression and function of KCC2, a neuroprotective ionic transporter that extrudes neuronal chloride. Importantly, no other transporter can rescue KCC2 deficit, and attenuated expression of KCC2 is strongly associated with circuit malfunction in chronic pain, epilepsy, neuro-degeneration, neuro-trauma, and other neuro-psychiatric illnesses. To isolate *Kcc2* gene expression-enhancing compounds, we screened 1057 cell growth-regulating compounds in cultured primary cortical neurons. We identified kenpaullone (KP), which enhanced *Kcc2/KCC2* expression and function in cultured rodent and human neurons by inhibiting GSK3ß. KP effectively reduced pathologic pain in preclinical mouse models of nerve constriction injury and bone cancer. In nerve-injury pain, KP restored *Kcc2* expression and GABA-evoked chloride reversal potential in the spinal cord dorsal horn. Delta-catenin, a phosphorylation-target of GSK3ß in neurons, activated the *Kcc2* promoter via Kaiso transcription factor. Validating this new pathway in-vivo, transient spinal over-expression of delta-catenin mimicked KP analgesia. With relevance for pathologic pain, our discoveries of a newly repurposed compound and a novel genetically-encoded mechanism that each enhance *Kcc2* gene expression enable us to re-normalize disrupted inhibitory neurotransmission through genetic re-programming.

## Introduction

In the mature vertebrate central nervous system (CNS), γ-aminobutyric acid (GABA) acts primarily as an inhibitory neurotransmitter and is critical for normal CNS functioning^1,2^. In chronic pain, GABA-ergic transmission is compromised, causing circuit malfunction and disrupting inhibitory networks^3–12^. Therefore, we recognize a mandate to discover new approaches for the restoration of physiologic GABA-ergic transmission. These innovations will increase our basic understanding of chronic, pathologic pain. Such discoveries will enable us to address the unmet medical need of chronic pain, with safer and more effective alternatives to opioids.

In the adult vertebrate CNS, the K^+^/Cl^−^ cotransporter KCC2 is expressed exclusively in neurons. KCC2 continuously extrudes chloride ions, thus ensuring that intracellular levels of chloride ions remain low as required for inhibitory GABA-ergic neurotransmission^13–19^. In chronic pathologic pain, KCC2 expression is attenuated in the primary sensory gate in spinal cord dorsal horn (SCDH) neurons. This key pathophysiological mechanism contributes to an excitation/inhibition imbalance because it corrupts inhibitory neurotransmission leading to inhibitory circuit malfunction^5,7,11,20–24^. Notably, there is no ‘back-up’ protein that can rescue the KCC2 expression deficit. Thus, we reasoned that if we could boost *Kcc2/KCC2* gene expression (*Kcc2* – rodent gene; *KCC2* – human gene), we could normalize inhibitory transmission for relief of chronic pain.

We therefore conducted an unbiased screen of cell growth-regulating compounds. We used such compounds because we assumed that a sizable number of them function by interfering with epigenetic and transcriptional machinery. Since mature neurons do not divide, these compounds are attractive candidates to upregulate gene expression of *Kcc2/KCC2*. This will in turn lower neuronal chloride levels. We identified such compounds by rigorous iterations of primary and secondary screening, and we selected one of them for in-depth exploration: kenpaullone (KP), a glycogen synthase kinase-3 (GSK3)/cyclin-dependent kinase (CDK) inhibitor^25,26^. We found that KP functioned as an analgesic in preclinical mouse models. Our data suggest that the cellular mechanism of action of KP in neurons was based on its GSK3ß-inhibitory function which enhanced *Kcc2/KCC2* gene expression. *Kcc2* up-regulation in turn relied on nuclear transfer of the neuronal catenin, δ-catenin (δ-cat)^27,28^. We found that in the nucleus δ-cat enhanced *Kcc2* gene expression via Kaiso transcription factors^29^. Increased *Kcc2* gene expression led to increased chloride extrusion by KCC2 transporter in neurons. We also documented attenuated *Kcc2* expression in the SCDH of mice with constriction nerve injury and rescue of *Kcc2* expression by KP. In these animals, pain-related behavior improved in response to KP. Importantly, patch-clamp recordings revealed more negative, thus electrically more stable GABA-evoked chloride reversal potentials in SCDH pain relay neurons after KP treatment.

## Results

### 1057 compound screen in primary cortical neurons for *Kcc2* gene expression enhancers identifies kenpaullone

To identify *Kcc2* gene expression enhancing compounds, we cultured primary cortical neurons from *Kcc2*-luciferase(LUC)-knockin (*Kcc2*-LUCki) mice^30,31^ and used LUC as readout for activity of the proximal *Kcc2* promoter (2.5kB^18^), which drives LUC in this transgenic mouse line. In our screening assay we recorded a Z’ factor of 0.94. We remain aware that this strategy will not select for long-range enhancers of *Kcc2* gene expression that act outside the 2.5kB core *Kcc2* promoter. We screened 1057 compounds, contained in two NCI libraries (Fig. 1, Suppl File S1), related to inhibition of growth of malignantly transformed cells. Iterative screening followed by measurements of *Kcc2* mRNA (RT-qPCR^18^) and intracellular chloride ([Cl-]i) (clomeleon chloride indicator protein^18,32^), led to decreasing numbers of re-confirmed hits, and we ended up with 4 “winner” compounds (Fig. 1). Of these we identified kenpaullone (KP), a GSK3/CDK kinase inhibitor, as a promising compound for further study based on its previous record of neuroprotection in translationally relevant preclinical models^33–36^.

**Fig. 1.**
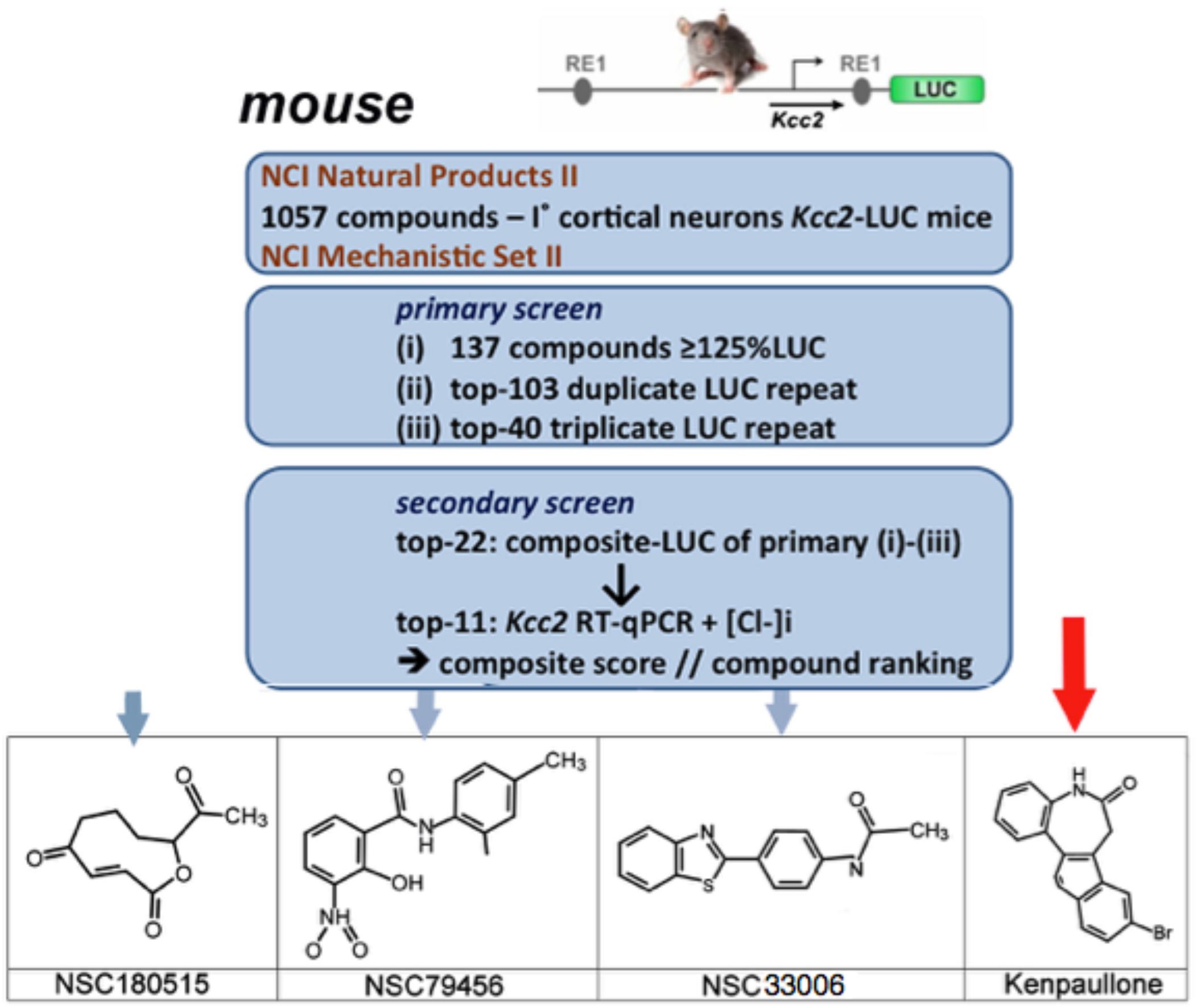
Compound screening for enhancers of Kcc2 expression in primary cortical neurons yields Kenpaullone. Screening paradigm using three rounds of primary screen based on luciferase (LUC) activity followed by secondary screen including *Kcc2* RT-qPCR and Clomeleon chloride imaging. Bottom panel: Four compound “winners” including Kenpaullone (KP); screening conducted in primary mouse neurons.

Our data establish that (i) KP enhanced *Kcc2* gene expression in rat and mouse primary cortical neurons (Fig. 2A, Suppl. File 1), (ii) this effect was dose-dependent when tested in rat neurons (Fig. 2A), (iii) in rat neurons, KP lowered [Cl-]i (Fig. 2B), (iv) this effect relied on chloride-extruding function of KCC2 transporter protein (Fig. 2C), (v) KP did not function as an enhancer of KCC2 transporter-mediated chloride efflux (Suppl Fig. 1), (vi) KP enhanced *Kcc2* gene expression and KCC2 chloride extrusion function with rather rapid kinetics within 3h (Suppl Fig. 1); (vii) in human primary cortical neurons, KP dose-dependently enhanced *KCC2* gene expression (Fig. 2D). The latter finding was accompanied by increased protein expression of KCC2 and synaptophysin (Fig. 2E-F), both of them co-localizing. We used synaptophysin as a marker of synaptic maturation and generally of a mature neuronal phenotype, which is rooted in increased expression of KCC2. These findings in rodent and human neurons indicate that our rationally designed screen identified a GSK3/CDK kinase inhibitor, KP, that enhanced *Kcc2/KCC2* gene expression and not KCC2-mediated chloride extrusion in CNS neurons. KP functioned as *Kcc2/KCC2* gene expression enhancer in mammals including humans. In human primary cortical neurons we found enhanced synaptic maturation and increased KCC2 expression, most likely causally linked.

**Fig. 2.**
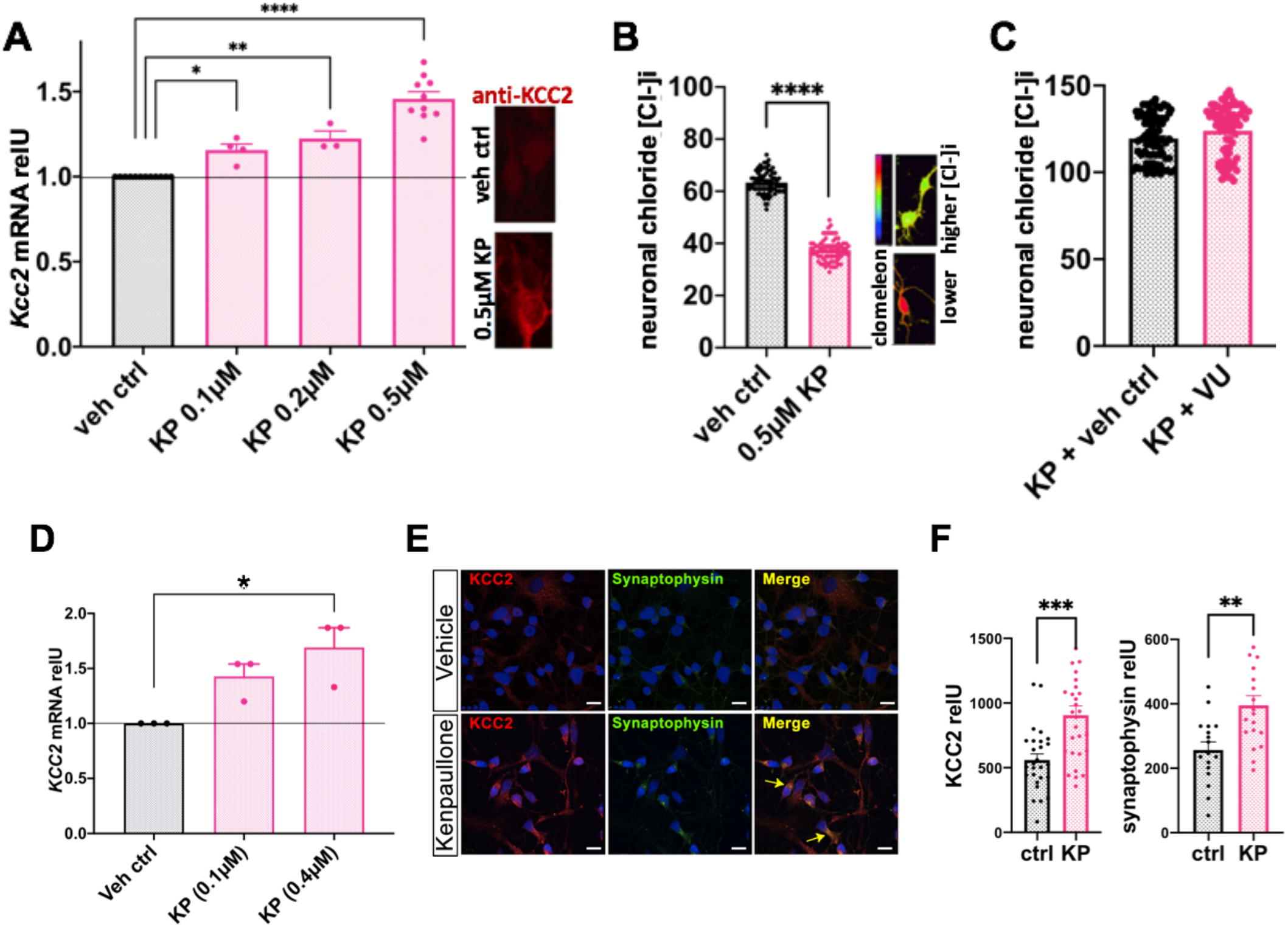
Kenpaullone enhances *Kcc2/KCC2* gene expression and function in rat and human primary cortical neurons. **A)** Primary rat cortical neurons. Bar diagram: dose-dependent increase in *Kcc2* mRNA expression after KP treatment, also reflected by increased protein expression as shown by KCC2 immuno-label (micrographs). Results represent the average mRNA expression of multiple independent neuronal cultures, n=12 (control), n=4 (0.1μM KP), n=3 (0.2μM KP), n=10 (0.5μM KP). *p<0.05, **p<0.01, ****p<0.0001, one-way ANOVA **B)** Primary rat cortical neurons. Neuronal [Cl-]i, measured with ratiometric chloride indicator, clomeleon, is robustly and significantly reduced after KP treatment. n≥75 neurons/ 3 independent cultures; **** p<0.0001, t-test **C)** Note that add-on treatment with KCC2-transport blocker, VU0240551 (2.5μM), leads to a [Cl-]i ≥120mM, for both, vehicle-treated and KP-treated, indicating that KP’s chloride lowering effect relies on KCC2 chloride extruding transport function. n≥75 neurons/ 3 independent cultures. **D)** Primary human cortical neurons. *KCC2* mRNA increases in a dose-dependent manner upon treatment with KP. Results represent the average mRNA expression of 3 independent neuronal cultures. *p<0.05, one-way ANOVA **E)** Primary human cortical neurons. Representative confocal images at DIV10 immuno-labelled for KCC2 and synaptic maturation-marker, synaptophysin, after vehicle or KP-treatment. Note co-localization (yellow arrow). Bar, 10μm. **F)** Primary human cortical neurons. Morphometry of immuno-cytochemistry shows increased KCC2 (62%; left-hand) and synaptophsin (54%; right-hand) expression after KP treatment, relative to vehicle. n=27-28 neurons analyzed for KCC2 \*\*\**p*<0.001; n=17-18 neurons analyzed for synaptophysin, \**p*<0.01, t-test.

### KP functions as an analgesic in-vivo

In view of these findings, we decided to address in-vivo analgesic effects of systemically applied KP. For this, we used two types of preclinical mouse models of pathologic pain: peripheral nerve constriction-induced neuropathic pain, using the PSNL method of nerve constriction injury^37,38^, and a bone cancer pain model that relies on implantation of mouse lung carcinoma cells into the marrow of the femur^39,40^. Nerve constriction injury, a widely used neuropathic pain model, was previously used to demonstrate down-regulated KCC2 expression in the spinal cord dorsal horn in pain^3^’^6^. Bone cancer pain is another clinically-relevant pain model. In particular, it fits the profile of KP, as the compound inhibits GSK3ß and CDKs, which might possibly be analgesic but also important for growth regulation of the implanted carcinoma cells.

We found that behavioral sensitization (mechanical allodynia) in both pain models, namely decrease in withdrawal thresholds in response to mechanical cues, was significantly reduced by KP treatment, as illustrated in Figs. 3A-B. Analgesic effects of KP were dose-dependent in both pain models. In nerve constriction, they were less accentuated at the lower dose of 10 mg/kg daily intraperitoneal (i.p) injections, and more pronounced at 30 mg/kg (Fig. 3A). In bone cancer pain, there was no significant analgesic effect at 10 mg/kg, and significant behavioral effects at 30 mg/kg (Fig. 3B). Importantly, we observed protracted analgesia, e.g. an almost complete elimination of pain hypersensitivity after nerve constriction on d7-14. The effects on bone cancer pain were also protracted and becoming apparent at d10 and d14, only at the higher dose of 30 mg/kg KP. This particular time-course can be interpreted as re-programming of sensitized nociception. Our findings are rather not suggestive of selective analgesic inhibition of a pro-algesic ion channel or receptor. Thus, in-vivo, KP functions as an analgesic when administered systemically, both in neuropathic pain model and in a bone cancer pain model. Of note and in contrast to its effective analgesic profile in bone cancer pain, KP did not significantly inhibit osteolysis/bone damage (Suppl. Fig. 3A), indicating that it was not containing growth of the implanted mouse lung carcinoma cells.

**Fig. 3.**
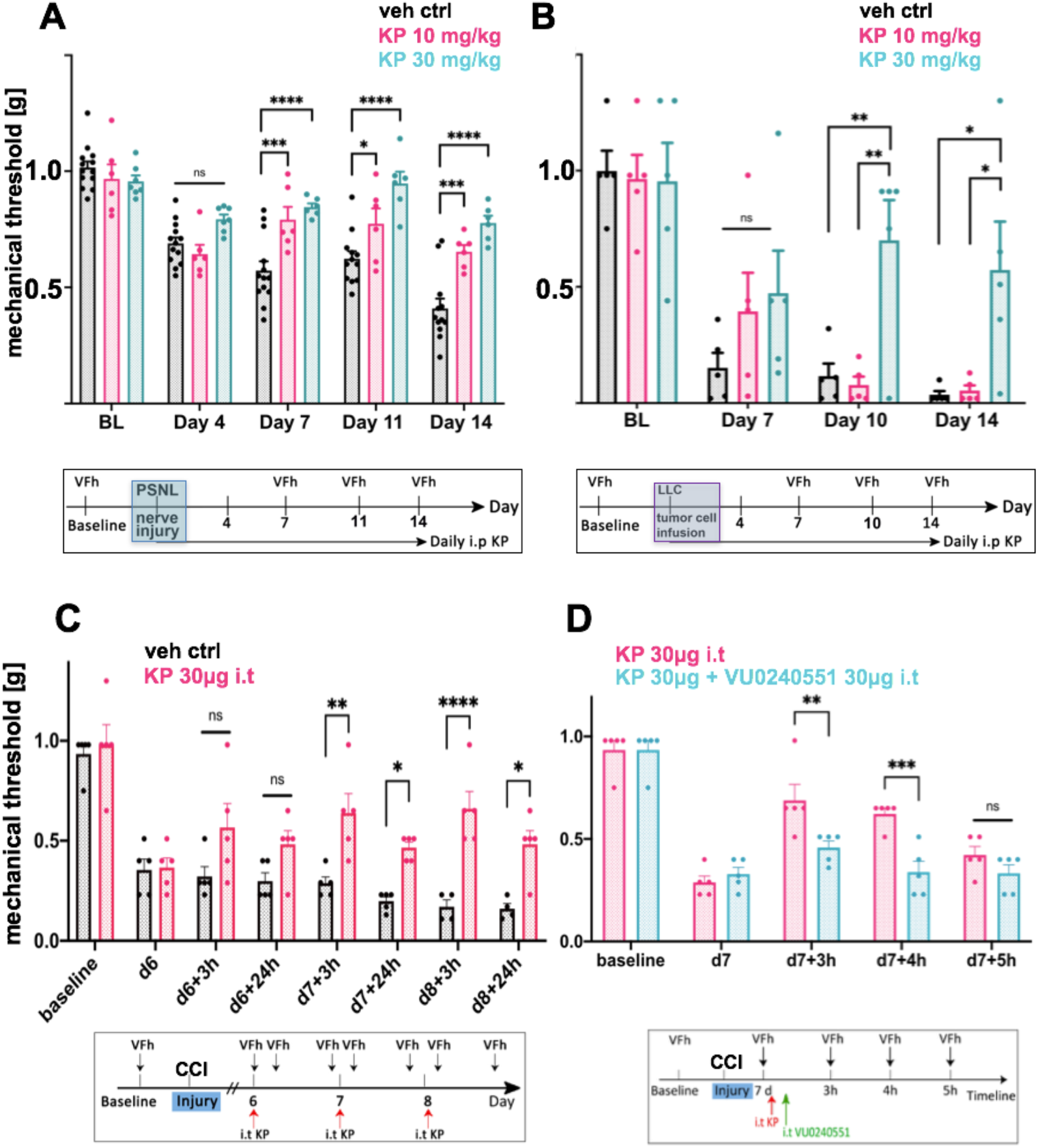
Kenpaullone is analgesic in mouse nerve constriction injury and bone cancer pain. **A)** Systemic KP is analgesic for nerve constriction injury pain. Bottom: Overview of timeline of systemic injection and behavioral metrics. Mice were injected intraperitoneally (i.p) with either 10mg/kg or 30mg/kg KP daily after nerve constriction injury (PSNL). Bar diagram: Analgesic effects of KP for sensitized mechanical withdrawal were dose-dependent, namely less accentuated at 10 mg/kg and more pronounced at 30 mg/kg. n=6-7 for KP treatment, n=13 for vehicle control; *p<0.05, ***p<0.001, ****p<0.0001, mixed model statistics **B)** Systemic KP reduces mechanical allodynia in bone cancer pain evoked by infusion of mouse LLC lung cancer cells into the femur. Bottom: overview of timeline of systemic injection and metrics. Mice were injected with KP as in A). Bars: analgesic effects of KP for sensitized mechanical withdrawal were significant at d10 and 14 for 30 mg/kg vs. 10 mg/kg and vs vehicle control. n=5 mice/group; *p<0.05, **p<0.01, two-way ANOVA **C)** Intrathecal (i.t) KP reduces mechanical allodynia in neuropathic pain following nerve constriction injury. Bottom: Overview of timeline of nerve constriction (CCI), i.t injection and behavioral metrics. Bar diagram: analgesic effects of KP for sensitized mechanical withdrawal were observed upon daily i.t injection (30μg), note increasing sensitization in vehicle injected controls. Withdrawal behavior was measured at two times after i.t injection, +3h, +24h. *p<0.05, n=5 mice/group, *p<0.05, **p<0.01, ****p<0.0001, two-way ANOVA **D)** i.t co-application of KP and specific KCC2 transport inhibitor compound, VU0240551 (30μg), blocks the central analgesic effects of KP. Behavior assays were conducted at 3h, 4h and 5 h on day 7 after i.t injection; n=5 mice/group; **p<0.01, ***p<0.001, two-way ANOVA

We then asked whether the analgesic effects of KP were mediated centrally at the spinal cord level. We injected KP intrathecally (i.t; 30 μg) and observed reduced mechanical allodynia in mice with nerve constriction injury, using the CCI method of nerve constriction (Fig 3C). The effect was apparent at 3h and persisted at 8h. This time-course is in keeping with the rapid effect on *Kcc2* gene regulation that we observed in primary cortical neurons (Suppl. Fig. 1). In order to learn whether the i.t analgesic effect of KP relies on KCC2 transporter-function, we next co-applied KP with the KCC2 chloride transport inhibitor VU0240551 and observed an elimination of the i.t analgesic effects of KP (Fig. 3D). This observation suggests that the central analgesic effect of KP depended on KCC2-mediated chloride extrusion.

In view of these beneficial effects of KP on pathologic pain, and to prepare for translation toward clinical use, we next addressed whether KP had undesirable effects on the CNS in terms of sedation, impairment of motor stamina, balance and coordination. Rotarod testing^41^ of KP-treated mice (10, 30 mg/kg; i.p.) showed that KP did not induce unwanted side effects (Suppl. Fig. 3B). We also assessed whether KP (30 mg/kg; i.p.) can trigger brain reward mechanisms by measuring conditioned place preference^42,43^, and there were no such effects (Suppl. Fig. 3C). Thus, KP functions as an analgesic in relevant preclinical mouse models and does not cause unwanted effects including effects on reward, sedation, lack of coordination, and reduced stamina. In pathologic pain, KP acts centrally to mediate analgesic effects, and this involves KCC2 chloride extrusion.

### KP renormalizes E_GABA_ in spinal cord dorsal horn by increasing *Kcc2* expression and function

With the above findings and in view of the SCDH as the likely site of analgesic action of KP^3,6^, we focused on SCDH *Kcc2* expression and function in nerve injury and response to KP. We found that in the SCDH, KP repaired attenuated *Kcc2* expression caused by partial sciatic nerve ligation (PSNL), at both the mRNA and protein levels (Fig. 4A-B). To measure *Kcc2* mRNA, we microdissected SCDH Rexed-layers I-II with laser capture followed by RT-qPCR (Fig. 4A). SCDH KCC2 protein expression was measured morphometrically after KCC2 immunolabeling using SCDH laminae I-II as the region-of-interest (Fig. 4B).

**Fig. 4.**
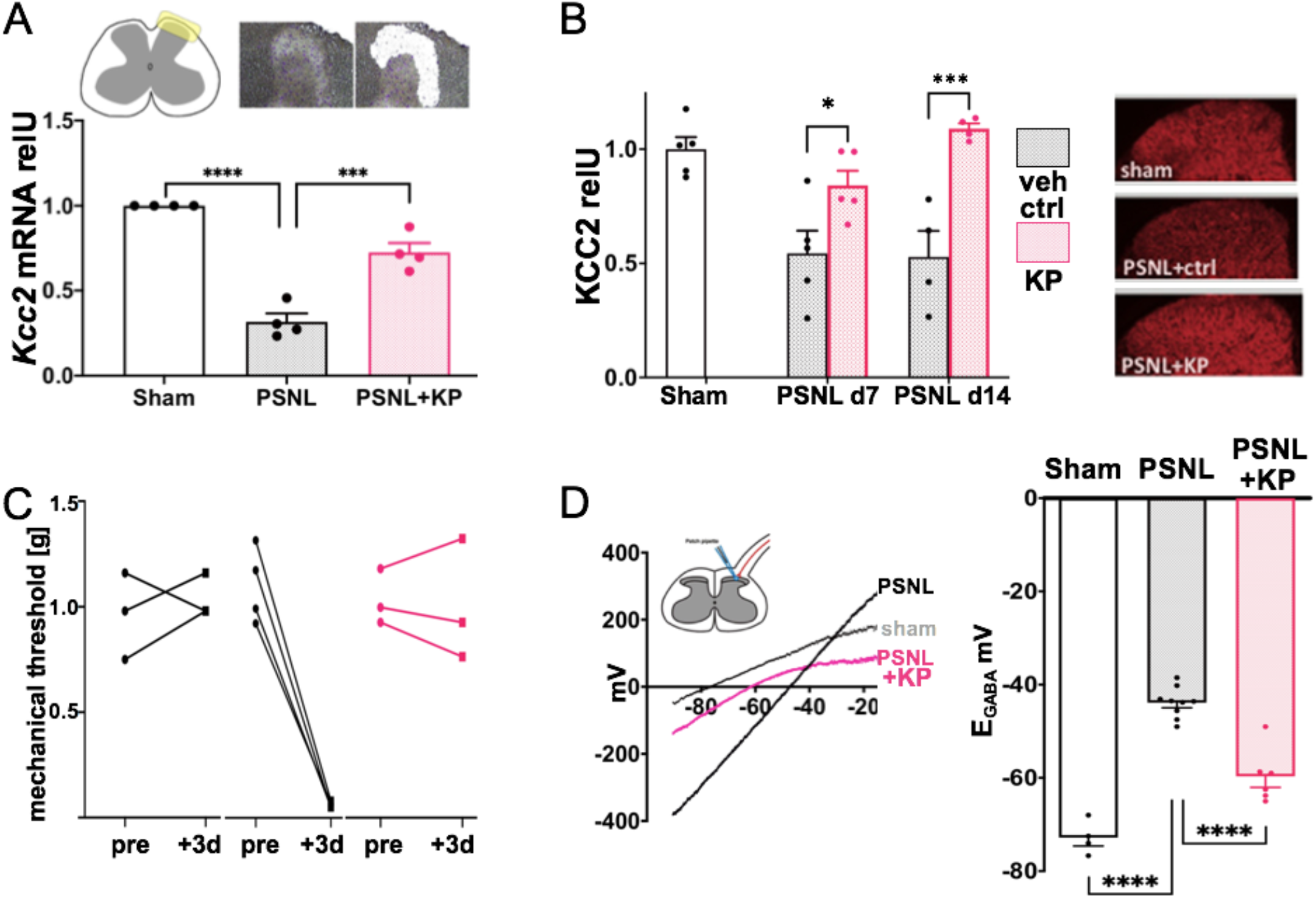
Kenpaullone re-normalizes E_GABA_ in spinal cord dorsal horn by increasing *Kcc2* expression/function. **A)** Top left: Lamina-I/II area of spinal cord dorsal horn (SCDH) is highlighted. Top right: image representation of lamina-I/II area before and after laser capture microdissection. Bottom, bar diagrams: KP (10mg/kg) rescues significantly attenuated *Kcc2* mRNA expression in the SCDH after nerve constriction injury (PSNL) vs vehicle treated mice; n=4 mice/group, **** p<0.0001, one-way ANOVA **B)** Bar diagrams: KP (10mg/kg) increases KCC2 protein expression in SCDH compared to vehicle control in PSNL nerve constriction injury. Right-hand panels: representative KCC2 immuno-staining of the SCDH in nerve constriction injury (PSNL). n=5 mice/group, *p<0.05; ***p<0.001, one-way ANOVA. **C)** Potent behavioral sensitization of juvenile mice after peripheral nerve constriction injury (PSNL) and its almost complete behavioral recovery after treatment with KP (30 mg/kg). n=3 mice sham, n=4 mice PSNL + vehicle, n= 3 mice PSNL + KP. Note accentuated responses for sensitization and rescue in younger mice vs adult mice (compare with Fig 3A). **D)** Spinal cord slices were isolated from the same juvenile mice as shown in panel C), and lamina-II neurons were examined for E_GABA_ using the perforated patch method, illustrated by the schematic, 1-3 neurons per mouse. The left-hand panel shows a representative *I–V* plot. Right-hand, bar diagram shows quantification of E_GABA_ indicating a significant depolarizing shift in sham vs PSNL with vehicle treatment (“PSNL”). Note significant hyperpolarization in response to KP treatment in PSNL mice. Of note, PSNL plus KP was not different from sham injury. n=4 neurons (sham), n=9 neurons (PSNL), n=6 neurons (PSNL+KP); \*\*\*\**p*<0.0001, one-way ANOVA

Nerve injury-induced downregulation of KCC2 has been shown to cause chloride reversal potential change in SCDH neurons after application of GABA (E_GABA_) in spinal cord slices^3^. We next investigated whether repair of attenuated *Kcc2* expression was associated with re-normalization of E_GABA_. Prior to interrogation of spinal cord slice preparations at 72h post-injury, we demonstrated robust mechanical allodynia of young mice by peripheral nerve constriction injury (PSNL) and its almost complete behavioral reversal by systemic treatment with KP at 30 mg/kg (Fig. 4C). We noted that PSNL injury sensitized the juvenile mice more than adults. Also, behavioral rescue with KP was very potent in juveniles. Next, in spinal cord slice preparations derived from these animals, SCDH lamina-II neurons were investigated by perforated patch-clamp (Fig. 4D, Suppl Fig 4). Our data show that the more positive, thus more excitable E_GABA_ reversal potential in PSNL mice was significantly lowered toward more negative values in KP-treated animals (Fig. 4D). Our documentation of re-normalization of E_GABA_ in SCDH lamina-II neurons is consistent with repair of *Kcc2* expression in SCDH in KP-treated mice. Thus, in a constriction nerve injury model in mice, the central analgesic action of KP relies on the enhanced gene expression of *Kcc2* and re-normalization of E_GABA_ in pain-relaying neurons in the SCDH.

### Cellular mechanism of action in neurons: GSK3B→δ-cat→Kaiso→*Kcc2*

These findings set up a compelling rationale to deconstruct the cellular mechanism of action of KP that accounts for its effects in SCDH neurons and thus its sensory effects. We conducted these studies in primary cortical neurons because 1) these neurons were used for our initial screen, 2) there was rodent-human similarity in terms of the effects of KP on *Kcc2/KCC2* gene expression, and 3) the effects of KP on *Kcc2* gene expression and KCC2 chloride transporter function in these neurons were highly similar to our findings in SCDH lamina-II neurons, which cannot be cultured as readily for mechanistic cellular studies.

First we addressed the question whether KP inhibiting GSK3ß or CDKs is responsible for increasing expression of *Kcc2.* Using a set of GSK3 inhibitors different from KP, we demonstrated increased expression of *Kcc2*. In contrast, a suite of CDK-inhibitory compounds did not increase (but decreased) expression of *Kcc2* (Fig. 5).

**Fig. 5.**
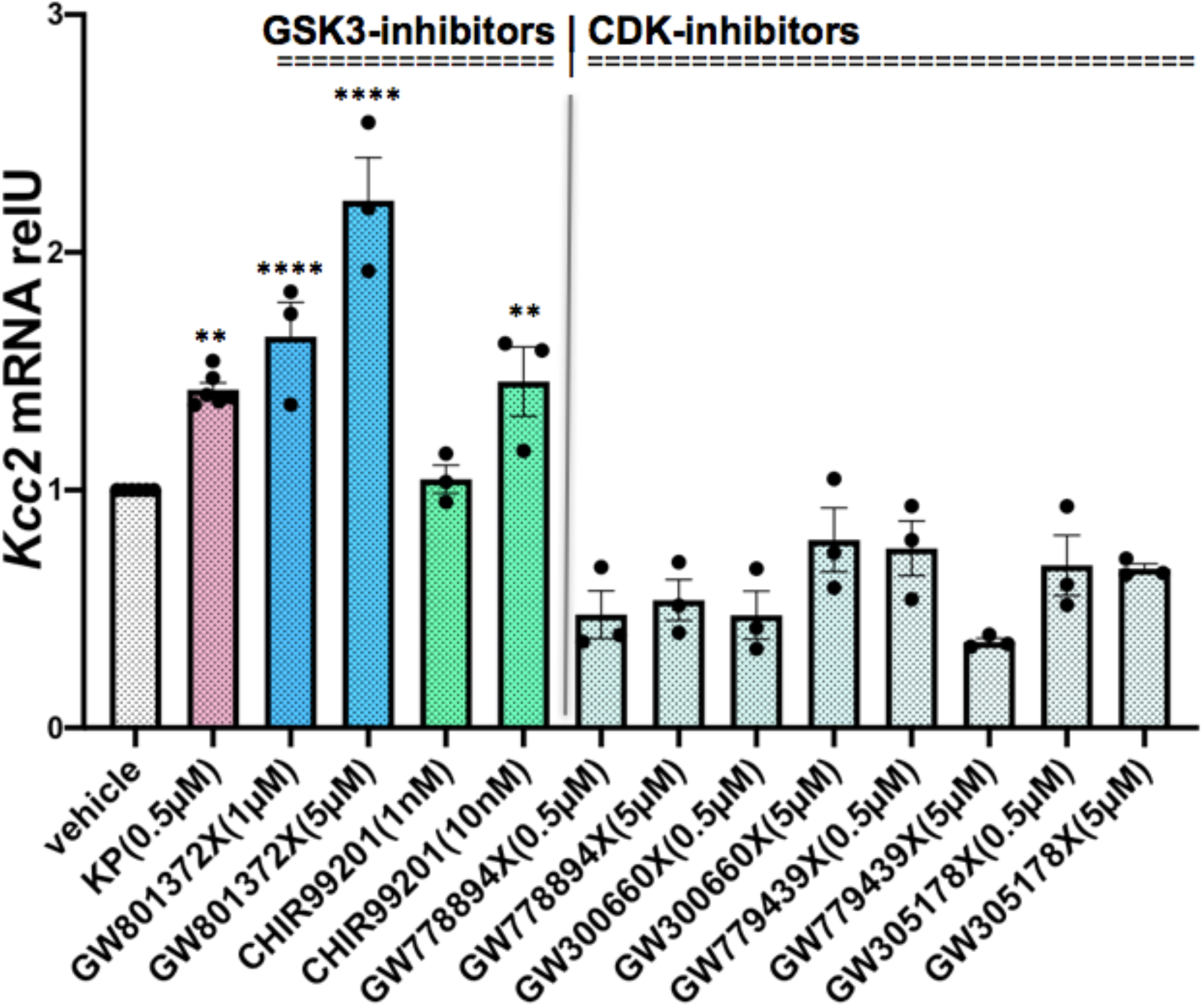
Kenpaullone increases Kcc2 expression in central neurons by inhibiting GSK3ß, not CDKs. GSK3-inhibitors increase *Kcc2* mRNA expression, measured by RT-qPCR, in a dose-dependent manner, whereas several CDK-inhibitors do not increase *Kcc2* mRNA expression, or even reduce it. Rat primary cortical neurons. Results represent the average mRNA expression of 3-6 independent neuronal cultures. **p<0.01, ****p<0.0001, compound vs vehicle, one-way ANOVA.

We then used a drug affinity responsive target stability (DARTS) assay^44^ to study direct binding of KP to GSK3ß (Fig. 6A, Suppl. File S2) in primary cortical neurons. Our affirmative findings are consistent with previous reports in cell lines^45^ yet novel in primary neurons. Of note, other known kinase targets of KP such as CDKs were not identified in our unbiased DARTS assay. This confirms our result with inhibitor compounds that it is the GSK3ß-inhibitory function of KP that is responsible for its *Kcc2/KCC2* gene expression enhancing property in neurons. We next sought to identify kinase targets of GSK3ß that are differentially phosphorylated in response to KP. Using unbiased phosphoproteomics assays in rat primary cortical neurons, we identified the neuronal catenin, δ-catenin (CTNND2; δ-cat)^46,47^. We found that the serine at position 259 was one site at which differential phosphorylation occurred short-term (1h) and persisted long-term (24h) in response to KP (Fig. 6B, Suppl File S3). The respective residue in human δ-cat is S276. ß-catenin (ß-cat) could also be a GSK3ß kinase target, yet ß-cat was not significantly differentially phosphorylated. δ-cat(S276) is found in phosphosite.org and has been previously described^48^, but its identification in primary neurons is novel. Catenin phosphorylation is known to facilitate its own intracellular degradation via ubiquitination^49,50^. To examine whether non-phosphorylated δ-cat traffics to the neuronal nucleus, we conducted specific δ-cat immunolabeling followed by confocal microscopy and morphometry. Our results indicate that KP treatment of primary neurons enhanced δ-cat nuclear transfer (Fig. 6C-D). Given the known binding of ß-cat/δ-cat^51^, we immunolabeled for ß-cat and obtained a similar result (Suppl Fig 6), thus corroborating the δ-cat nuclear transfer results.

**Fig. 6.**
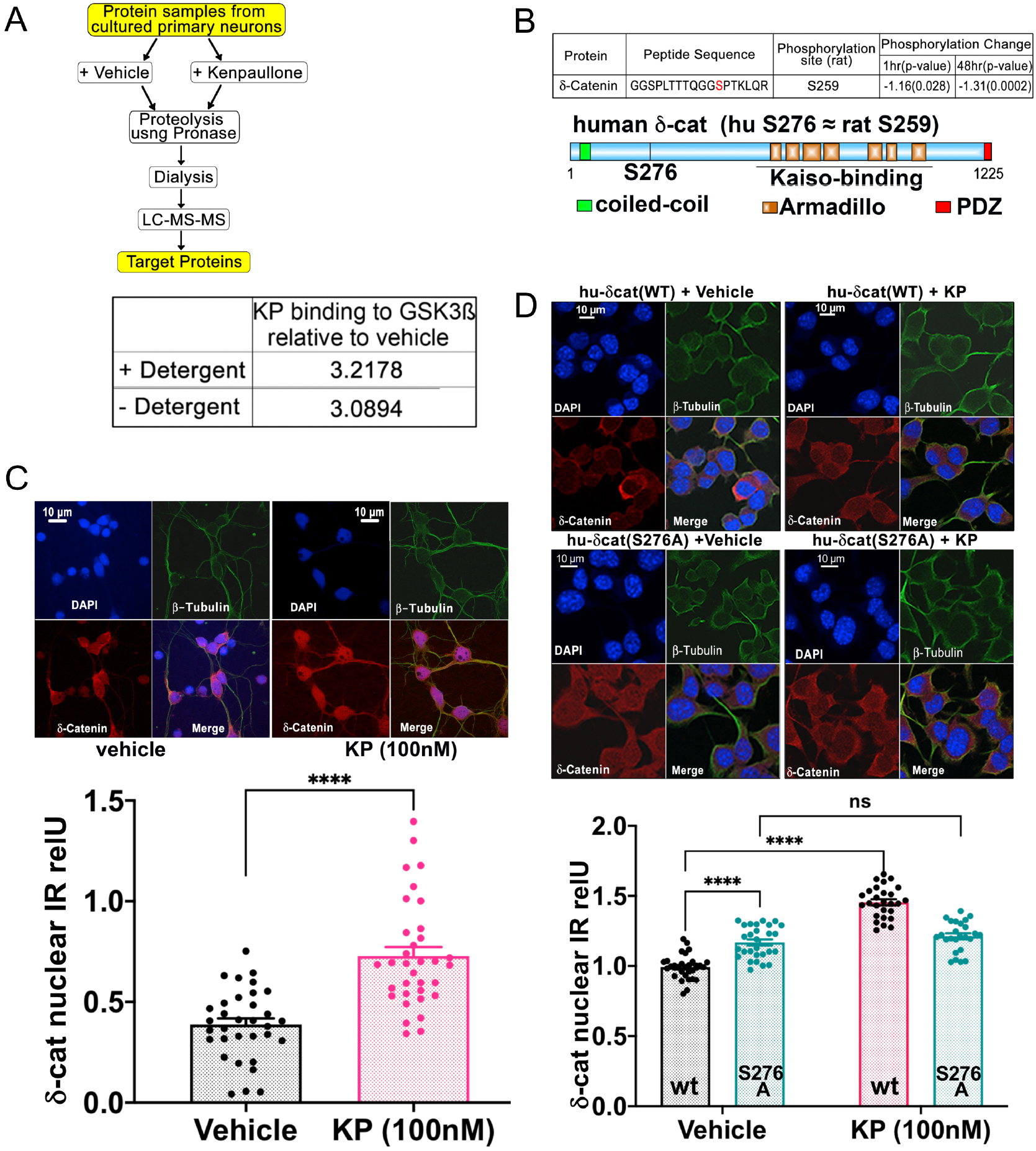
Cellular mechanism of action of Kenpaullone in central neurons. **A)** Left panel: DARTS methodology to identify proteins that bind to KP in rat primary cortical neurons. Right panel: KP binding to GSK3β is independent of detergent treatment of the protein sample preparation from the neuronal culture. Of note, binding to GSK3ß was documented whereas binding to CDKs was not (see Suppl dataset File S2 DARTS). **B)** Top panel: Phosphoproteomics assays reveal S259 phosphorylation target in δ-cat protein after KP treatment of rat primary cortical neurons, significant de-phosphorylation resulted after 1h treatment and was sustained at 24h (see Suppl dataset phosphoproteomics). Bottom: schematic representation of structure of human δ-cat (CTNND2) showing functional domains. Human residue S276 matches rat S259. The Armadillo domain region plays a key role in transcription factor Kaiso binding to δ-cat. **C)** Top panel: Representative immuno-labeling of ß_III_-tubulin (neuronal isoform of ß-tubulin) and δ-cat before and after KP treatment in rat primary cortical neurons, DAPI counterstain for nuclei. Bottom bar diagram: KP significantly increases δ-cat nuclear translocation. n=33-34 neurons/group,****p<0.001 KP vs vehicle, t-test. **D)** Top panel: Representative immuno-labeling of ß_III_-tubulin and δ-cat before and after KP treatment in differentiated N2a mouse neural cells which were transfected with either hu-δ-cat(WT), hu-δ-cat(S276A) or control vector. Bottom Panel: KP significantly enhances nuclear transfer of δ-cat when transfected with δ-cat(WT). Mutation δ-cat(S276A) increases nuclear transfer, but treatment with KP had no effect. n=25-30 neurons/group ****p<0.0001 vs vehicle, mixed effects statistics.

To investigate the relevance of δ-cat residue S276, we switched to N2a neural cells because these cells transfect at higher efficiency than primary cortical neurons. In our cultures, N2a cells expressed neuronal ß_III_-tubulin in elongated processes (Fig. 6E, Suppl Fig 6) as an indicator of their neuronal differentiation. Furthermore, nuclear transfer of δ-cat was significantly enhanced upon KP treatment in N2a cells transfected with human δ-cat(WT) (Fig. 6E-F). This increase in nuclear transfer was very similar to the trafficking we recorded in primary cortical neurons, thus validating the cell line. To determine the function of δ-cat residue S276 we then transfected δ-cat(S276A), which resulted in significantly increased nuclear transfer of δ-cat(S276A), using morphometry. Of note, there was no increase of nuclear transfer of δ-cat(S276A) upon KP treatment (Fig. 6E-F).

Thus, δ-cat(S259/S276) (rat/human) is very likely a relevant phosphorylation site in δ-cat and a GSK3ß kinase target in neurons. Inhibition of GSK3ß or rendering S276 phosphorylation-resistant enhances nuclear transfer of δ-cat, also of its binding partner ß-cat.

We conclude that catenins enhance *Kcc2* gene expression in neurons. Consequently, we explored their effects on the *Kcc2* promoter.

We identified two Kaiso binding sites (potential sites for δ-cat^29,52^) in the *Kcc2* proximal promoter using computational methods^53^. Two Kaiso binding sites bracketed the transcriptional start site (TSS) of the *Kcc2* gene (Fig. 7A). In rat primary cortical neurons, δ-cat was bound to both Kaiso sites in the *Kcc2* promoter (Fig. 7A). Interestingly, treatment with KP inhibited the binding of δ-cat to the upstream site. Binding of δ-cat to the site 3’ to the TSS was enhanced, suggesting that the upstream site functions as repressor and the 3’ site as enhancer. ß-cat bound to a TCF (T-cell factor) DNA-binding site^54^ close to the 5’ RE-1 site within the *Kcc2* promoter (Suppl Fig 7)^18^, and treatment of cells with KP significantly increased this interaction, indicative of enhancement.

**Fig. 7.**
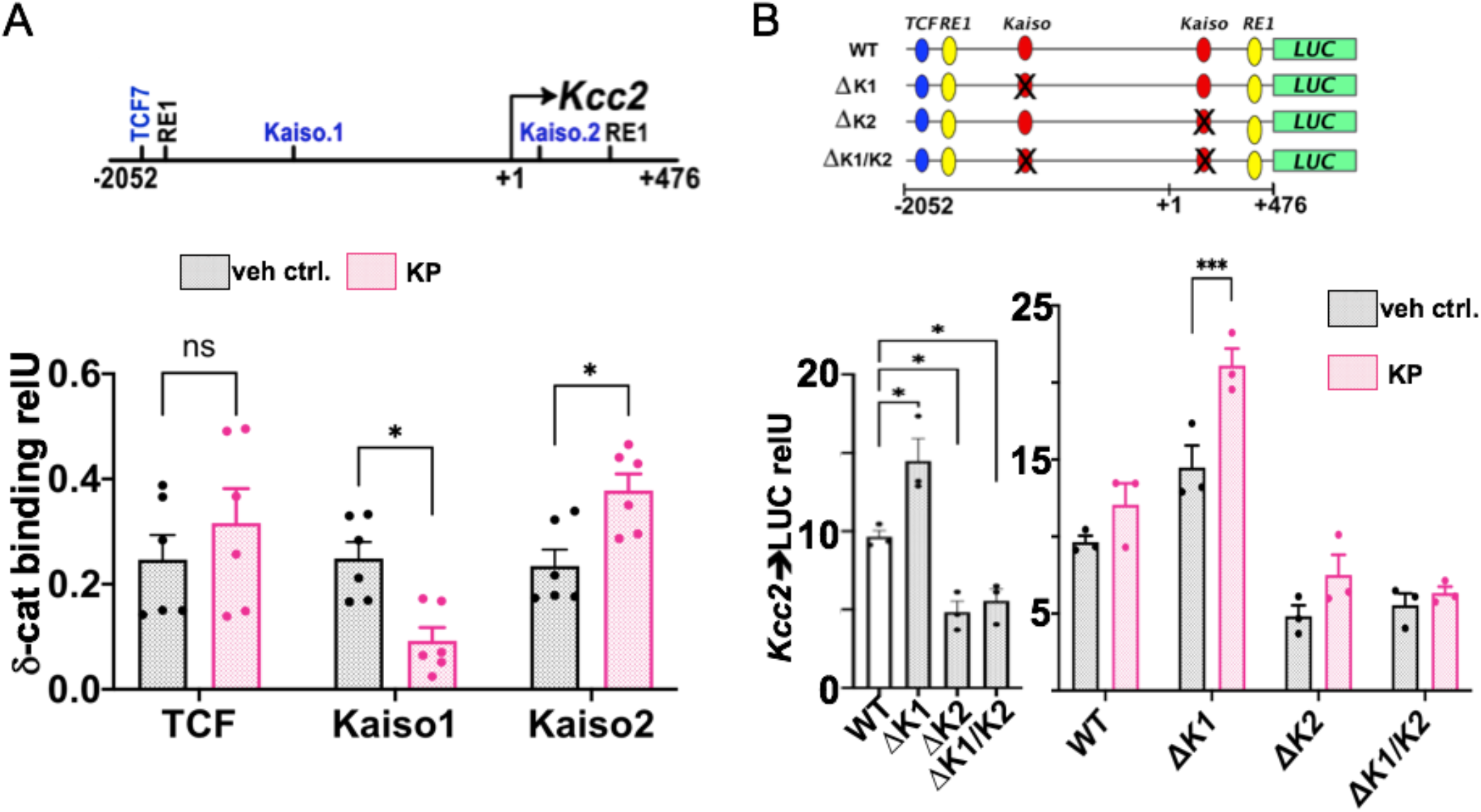
Kenpaullone regulates *Kcc2* promoter activity via δ-cat and two Kaiso sites. **A)** Structure of mouse *Kcc2* gene encompassing 2.5kb surrounding the transcription start site (TSS; +1). Location of DNA binding sites: Kaiso1 (−1456 to −1449), Kaiso2 (+83 to +90) and TCF (−1845 to −1838) relative to TSS, all three sites can bind δ-cat via Kaiso (Kaiso1, 2 sites) and ß-cat (TCF). Bottom, bar diagram: Chromatin immuno-precipitation (ChIP) using anti-δ-cat antibody in rat primary cortical neurons reveals binding of δ-cat to all three sites. KP treatment significantly increases binding of δ-cat to the *Kcc2* promoter on the Kaiso2 binding site, significantly reduced binding to Kaiso1, and non-significant increase at TCF. n=6 independent neuronal cultures were subjected to ChIP; *p<0.05 KP-treatment vs vehicle, t-test. **B)** Top panel: mouse *Kcc2* promoter constructs, Kaiso1, −2 were deleted and a ΔK1/K2 construct was built devoid of both sites. Dual-RE1 sites and TCF site shown for orientation, also TSS at +1. Bottom, bar diagrams: Luciferase (LUC) activity of *Kcc2* promoter constructs in N2a cells with neuronal differentiation (Suppl Fig. 6). Left-hand: ΔK1 significantly increased over WT, ΔK2 and ΔK1/K2 significantly decreased vs WT, n=3 independent culture per transfection, *p<0.05, 1-way ANOVA. Right-hand: repressive Kaiso1 binding site deletion with enhanced activity of the *Kcc2* promoter, significantly enhanced further in response to KP. n=3 independent cultures per transfection; ***p<0.001 KP-treatment vs vehicle for the respective construct, t-test.

We then built promoter expression constructs with rationally-targeted deletions to interrogate the effects of the Kaiso- and TCF-binding sites on activity of the *Kcc2* promoter and to determine if this activity was regulated by KP. For ease of transfection, key to this method, we again used N2a neural cells. The 5’ and 3’ δ-cat Kaiso binding sites functioned in repressive and enhancing manners, respectively (Fig. 7B). Presence of the 3’ δ-cat Kaiso binding site and absence of the 5’ site led to significantly enhanced activity of the *Kcc2* promoter upon treatment with KP. Deletion of both Kaiso sites led to markedly reduced promoter activity and non-responsiveness of the construct to KP. Deletion of the TCF binding-site from the *Kcc2* promoter did not change *Kcc2* promoter activity or its response to KP treatment (Suppl Fig. 7). However, the triple-deletion of both Kaiso sites and the TCF site rendered the construct minimally active and completely non-responsive to KP.

These data suggest that δ-cat, a kinase target of GSK3ß in CNS neurons as we show, traffics to the nucleus increasingly upon GSK3ß inhibition. In the nucleus, δ-cat interacts with the *Kcc2* promoter to enhance *Kcc2* expression via two Kaiso DNA-binding sites. ß-cat co-traffics to the nucleus with δ-cat in response to KP. ß-cat is not a significant neuronal GSK3ß kinase target (Suppl File S3), and plays an ancillary role in enhancement of *Kcc2* gene expression.

### δ-cat spinal transgenesis is analgesic in nerve constriction injury

We hypothesized that δ-cat, when expressed as a spinal transgene in sensory relay neurons, will facilitate analgesia in nerve constriction injury. We first documented that a δ-cat transgene increases *Kcc2* expression in N2a neural cells, and that *Kcc2* expression levels were slightly elevated when using δ-cat(S276A) (Fig. 8A). Thus, human δ-cat transgenes mimic the effects of KP in a mouse neural cell line.

**Fig. 8.**
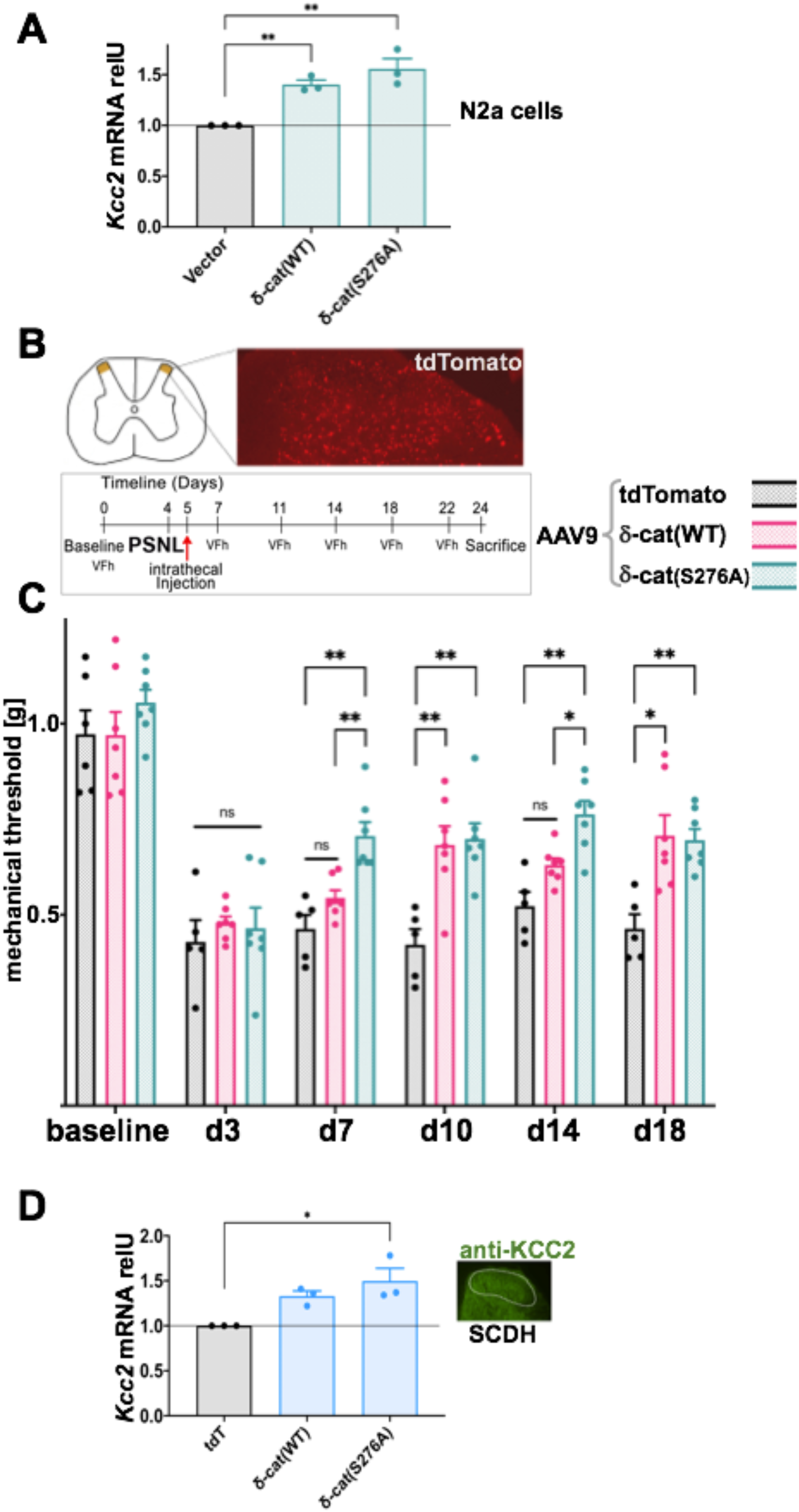
δ-cat spinal transgenesis is analgesic in nerve constriction injury. **A)** δ-cat(WT) transgene significantly increases *Kcc2* mRNA expression in differentiated N2a cells, and expression level was slightly elevated when transfecting δ-cat(S276A), both significantly increased over WT. n=3 independent cultures were subjected to transfection with the δ-cat constructs; **p<0.01 δ-cat construct vs control transfection, one-way ANOVA. **B)** Top, schematic and micrograph: SCDH showing lamina-I/II (left) and td-Tomato expression in SCDH neurons 3d post-i.t injection of AAV9-tdTomato; below - schematic: Timeline for behavioral testing after constriction nerve injury and subsequent i.t injection of AAV9 transgenesis vectors. **C)** Mechanical withdrawal thresholds after nerve constriction injury (PSNL). Significant improvement of sensitization for both δ-cat(WT) and δ-cat(S276A) constructs with sustained benefit over 2 weeks. Note more potent analgesia by δ-cat(S276A) based on the data on d7, d14. n=5-6 mice/tdTomato ctrl, n=7 mice/δ-cat transgenes; *p<0.05 **p<0.01, mixed effects statistics. **D)** *Kcc2* mRNA in microdissected SCDH was significantly increased in δ-cat(S276A) as assessed by RT-qPCR. For δ-cat(WT), *Kcc2* abundance was elevated but not to significant levels likely because of viral transduction of only a fraction of sensory relay neurons in the SCDH. n=3 mice/group, * p<0.05 δ-cat(S276A) vs tdTomato control, one-way ANOVA.

We then constructed AAV9 vectors harboring human δ-cat and δ-cat(S276A), driven by the minimal human neuronal synapsin promoter (huSyn^55^). We used AAV9 and huSyn because in a previous in-depth study, AAV9 harboring fluorescent reporter driven by huSyn, upon i.t injection, readily transduced spinal neurons and spared DRG primary afferent neurons^56^. We used synapsin-tdTomato as control and injected 5×10^9^ viral genomes (5 μL; i.t) of each construct. Assessment of tdTomato fluorescence 3d post injection revealed spinal transgenesis that was evenly manifesting in the SCDH (Fig 8B), in keeping with the above-mentioned previous study^56^. We then measured mechanical withdrawal thresholds after nerve constriction injury (PSNL) (Fig 8B-C). We found significantly less sensitization for δ-cat(S276A), with a protracted start of effect on d7 post-injection, and a sustained benefit until the end of experiment at δ-18 (Fig. 8C). δ-cat(S276A) showed increased effect vs. δ-cat(WT) on d7 and d14. δ-cat(WT) was significantly effective on d11, d18. In accordance with behavioral findings, *Kcc2* mRNA in microdissected SCDH was significantly increased in δ-cat(S276A) (Fig 8D). For δ-cat(WT), *Kcc2* mRNA abundance was elevated but not to significant levels likely because of viral transduction of only a fraction of sensory relay neurons in the SCDH. Thus, δ-cat(S276A) spinal transgenesis via AAV9 is sufficient to evoke analgesia after nerve constriction injury, to lesser degree also with δ-cat(WT). Importantly, δ-cat(S276A) spinal transgenesis was accompanied by increased expression of *Kcc2* in the SCDH. These findings suggest that our proposed mechanism of *Kcc2* gene expression enhancement by KP, via nuclear action of δ-cat at the *Kcc2* promoter, which we established in primary neurons and neural cells, remained valid in a nerve constriction preclinical pain model in mice. Moreover, our discovery points toward a genetically-encoded approach that can be developed for translation into clinical use as an alternative or complement to KP-related small molecules.

## Discssion

Evidence that reduced expression of the neuronal chloride extruding transporter, KCC2, critically contributes to chronic pathologic pain led us to seek a small molecule which increased its gene expression. Using measures of *Kcc2* promoter activity, *Kcc2* mRNA abundance, and [Cl-]i, we conducted an unbiased screen in primary cortical neurons of two NCI libraries containing 1057 compounds that inhibit growth of malignantly-transformed cells. We reasoned that a sizable number of these compounds would interfere with epigenetic and gene expression machinery, which renders them suitable candidates to function as *Kcc2/KCC2* expression enhancers in non-dividing neurons. Our screen identified kenpaullone (KP), a GSK3/CDK kinase inhibitor with advantageous neuroprotective properties^25,26,33–36^. Our studies of KP revealed the following principal findings: 1) KP directly enhances *Kcc2/KCC2* gene expression (not KCC2 transporter function) in a concentration-dependent manner and lowers [Cl-]i in cultured mouse, rat, and human neurons; 2) Systemic administration of KP to mice attenuates constriction nerve injury pain and bone cancer pain in preclinical models in a dose-dependent manner; 3) Intrathecal administration of KP to mice attenuates constriction nerve injury pain depending on spinal KCC2 chloride transporter activity; 4) Systemic administration of KP to mice with constriction nerve injury repairs defective *Kcc2* gene expression in SCDH neurons and shifts the GABA-evoked chloride reversal potential to more negative and electrically stable measures; 5) The mechanism by which KP enhances *Kcc2* gene expression is by binding to and inhibiting GSK3ß, inhibiting phosphorylation of δ2-cat at position S259 in rat (=276 in human), which increases nuclear transfer of δ2-cat. In the nucleus, δ2-cat binds to and enhances the *Kcc2* promoter via two Kaiso-binding sites surrounding the TSS of the *Kcc2* gene. 6) Testing whether this mechanism that we elucidated in neural cells is valid in live animals, we establish that spinal transgenesis of δ2-cat(S276A) attenuates constriction nerve injury pain in mice. We conclude that KP and the new GSK3ß→δ2-cat→Kaiso→*Kcc2* signaling pathway may represent a strategic bridge-head for therapeutics development for treatment of pathologic pain. Beyond pain, this could also apply to other neurologic and mental health conditions in which restoration of KCC2 function is important, such as epilepsy, traumatic spinal cord and brain injury, neurodegeneration and neurodevelopmental disorders. Our proposed analgesic mechanism is summarized in Fig. 9.

**Fig. 9.**
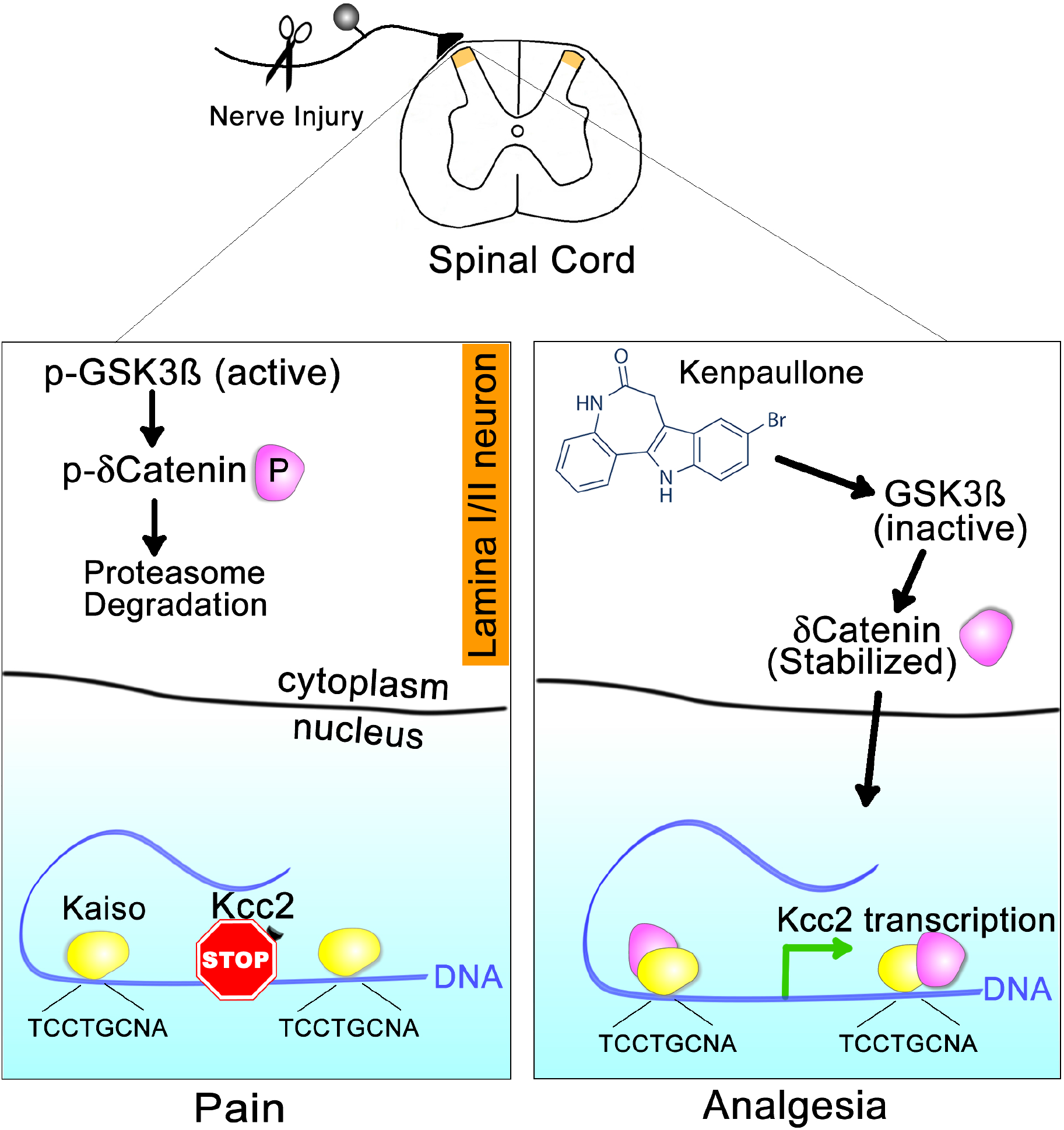
Analgesic mechanism of action of Kenpaullone. Left panel: Nerve injury facilitates activation of GSK3β kinase in pain relay neurons in layer-II of the SCDH. GSK3ß phosphorylates δ-cat in the cytoplasm at S259/S276. Phospho-δ-cat is unstable, undergoes ubiquitination and subsequent degradation. In the nucleus, Kaiso recruits repressive transcription factors on the *Kcc2* gene promoter which leads to overall repressed *Kcc2* transcription. Right panel: Treatment with KP inactivates GSK3β in SCDH layer-II pain relay neurons. In turn, this leads to de-phosphorylation and stabilization of δ-cat which enters the nucleus. In the nucleus, δ-cat binds to Kaiso transcription factor complex and displaces repressive transcription factors. This leads to a net enhancement of *Kcc2* expression.

We identified compounds from two NCI libraries that can interfere with gene regulation in CNS neurons. Primary developing cortical neurons enabled us to identify compounds that can enhance the *Kcc2* promoter, and we selected KP as the “winner” compound for in-depth exploration. Though KP is predicted to have multiple targets in CNS neurons, we provide evidence that KP binds to neuronal GSK3ß and not to CDKs. In addition, it is known that KP inhibits GSK3ß with highest potency from amongst known targets^25,45,57^. We also present evidence that GSK3ß inhibition by KP directly upregulates *Kcc2* gene expression via the δ-catenin-Kaiso pathway. This mechanism of a GSK3-inhibitory compound has not been reported previously. Of note, this novel concept held true in primary human neurons in which KP also enhanced synaptic maturation. The respective marker, synaptophysin, showed remarkable co-localization with upregulated KCC2 protein. We remain aware that KP can inhibit other kinases, but our data suggest that inhibition of GSK3ß and subsequent enhancement of *Kcc2* gene expression via δ-catenin are very important, perhaps dominant mechanisms of action of KP as it attenuates pathologic pain. Additionally, δ-cat-Kaiso likely affects multiple neuronal genes, but our data suggest that enhanced *Kcc2* gene expression and KCC2 function are the major analgesic effector mechanisms of KP. Another argument, mechanistically weaker but translationally relevant, is the absence of unwanted effects of our *Kcc2* expression-enhancing strategy on choice behavior, motor stamina, and coordination. Effective targeting of multiple pathways would likely impact these behaviors. However, the behavioral profile for KP was similarly benign as that of KCC2 chloride-extrusion enhancing compounds^6^.

Our screening strategy allowed us to address a fundamental problem of pathological pain. This vastly unmet medical need, rooted in its chronicity and overall debilitating impact, is driven by genetic reprogramming, which results in a maladaptive phenotype^5,9,12,58,59^. One very important mechanism to contribute to the maladaptive phenotype is attenuated expression of *Kcc2* because of its relevance for inhibitory transmission in pain-relevant neural circuits^3,6,7,20,21,23^. This has also been postulated in other pathologic conditions of the CNS as a general pathogenic feature^60,61^. Regulation of *Kcc2* gene expression by GSK3ß and its kinase target δ-cat is a novel insight of our study. This concept will permit rational exploration of links between GSK3ß→δ-cat and attenuated *Kcc2/KCC2* gene expression and the resulting malfunction of inhibitory neurotransmission in several other relevant neurologic and psychiatric conditions,such as epilepsy, traumatic brain/spinal cord injury, Rett Syndrome, Autism Spectrum Disorders, and perhaps also Alzheimer’s Disease and other neurodegenerative diseases^62–70^. We selected KP because of its previously reported neuroprotective properties for spinal motoneurons, brainstem auditory relay neurons, and hypoxia-injured hippocampal neurons^33–36,71,72^. It is possible that there was a unifying mechanism of *Kcc2* expression enhancement by KP in these previous studies. Neuroprotective properties for a novel analgesic are welcome because chronic pain is associated with non-resolving neural injury mediated by neuroinflammation^73^. The repurposing of a GSK3ß-inhibitory compound as an analgesic reprogramming agent that upregulates *Kcc2/KCC2* expression links chronic pathologic pain to neurodegeneration, at both the basic science and translational neuroscience levels.

Our work characterizes a novel strategy to treat chronic pathologic pain and also increases our basic understanding of pain and sensory transduction. With a focus on enhanced expression of *Kcc2* in central neurons as an analgesic strategy, we repurpose a compound as a genomic reprogramming agent that reverts the expression of a key dysregulated gene. This puts us in position to address a mechanism that underlies neural malplasticity in chronic pain: ineffective inhibitory neurotransmission in pain-relevant neural circuits caused by genomic dysregulation^9,58,59^, such as the lack of expression of *Kcc2*^3,6,7,20,21,23^. Of practical/translational importance, we found that KP is non-sedative and does not affect motor stamina, coordination, or choice behavior. Furthermore, we target-validated KP in human primary neurons. In aggregate, we identified a repurposed compound, KP, and a genetically-encoded cellular signaling pathway, GSK3ß→δ-cat→Kaiso→*Kcc2*, which were not previously known to up-regulate *Kcc2.* Both of these *Kcc2*-enhancing approaches can be developed translationally into clinical neuroscience applications.

Our finding that KP acts as an analgesic in a bone cancer pain model is novel and noteworthy, particularly in the arena of cancer pain. This finding requires additional research to further elucidate the underlying cellular and neural circuit mechanisms, e.g in the SCDH. Interestingly, effective analgesia was observed at the higher KP dose whereas bone lesions caused by implanted cancer cells were not significantly affected. However, given the important established roles of CDKs and GSK3 in growth-regulation of cancer cells, future studies will elucidate whether KP can enhance co-applied cell-ablative strategies, and whether higher doses of KP might improve analgesia and contain cancer cell growth.

Our approach as described here for targeting of pain is innovative. In the context of regulating neuronal chloride in the SCDH for analgesia, the previous discovery of small molecules that enhance KCC2 chloride extrusion is noteworthy^6^. This latter study brought full circle the initial landmark observation of lack of expression of KCC2 in SCDH neurons after peripheral injury^3^. Together, these previous studies demonstrate causality of KCC2 downregulation in pathologic pain. Interestingly, recent results suggested that this mechanism also might apply to human spinal pathophysiology in pathologic pain^22^.

In an elegant and very recently published study, small molecules were selected to enhance expression of *Kcc2/KCC2* for effective rescue of a modeled Rett Syndrome phenotype^74^.

It is our view that enhanced *Kcc2/KCC2* gene expression, based on KP treatment or δ-cat transgenesis as presented here, will complement direct enhancement of KCC2 chloride extrusion in targeting pathologic pain. Complementary use will help overcome recalcitrant lack of expression and function of KCC2 in pain relay neurons, as might be expected in clinical cases of “refractory” chronic pain. Clinical combination-use of *KCC2* expression enhancers with analgesic compounds that have different mechanisms of action will be advantageous because renormalized inhibitory transmission will likely cause improved effectiveness of other compounds, such as gabapentinoids. Supporting this concept, a very recent study reported effective analgesia of GABAA receptor a2/a3 agonists when enhancing KCC2 extrusion in neuropathic pain^75^.

## Supporting information

Supplemental data sheet 1

Supplemental data sheet 2

Supplemental dat a sheet 3

## Acknowledgements

Invaluable comments on the manuscript were provided by Duke University colleagues Drs. James McNamara, Albert LaSpada, Nicole Calakos, Rochelle Schwartz-Bloom and Sidney Simon.

Viral DNA was packaged by the Duke Neurotransgenesis Viral Vector Core, lead-scientist Dr. Boris Kantor. Phospho-proteomics and DARTS analysis were conducted with the expert help of the Duke Proteomics Core Facility, Director Dr. M. Arthur Mosely and lead-scientist Dr. Erik Soderblom.

## Author contributions

conducted experiments - MY, YC, CYJ, GC, KW, ML, QZ, ZLW, WL; conceptual input - MY, YC, AB, JB, RRJ, WL; wrote paper - MY, YC, CJ, GC, ML, JB, RRJ, WL

## Data and materials availability

All data associated with this study are available in the main text or the supplementary materials.

## Methods

### Screening in primary cortical neurons from *Kcc2*-LUC transgenic mice, rat primary cortical neurons

Transgenic mice that express red-shifted luciferase (LUC) under the control of the *Kcc2* promoter (−2052/+476, described in^18^), inserted into the Rosa26 locus, were previously described by us^30,31^. We generated primary cortical neuronal cultures from newborn (p0) mice of this line^30,31^.

After one week in culture, neurons were treated with compounds at 100 nM for 48h; LUC activity was determined for each compound with vehicle and trichostatin-A as negative and positive controls^18^.

Experimental flow is depicted in Fig. 1A.

Rat primary cortical neuronal cultures were maintained as in^18^.

### Compound libraries

NCI compound libraries Natural Products II and Mechanistic Diversity Set II were obtained from NCI. 1057 compounds are listed in Supplemental Table 1. See also: https://dtp.cancer.gov/organization/dscb/obtaining/available_plates.htm

### Human neuronal cultures

The neuron-enriched cultures were established from male and female human fetal cortical specimens at 15 – 20 weeks of gestation. The protocols for tissue processing complied with all federal and institutional guidelines. Human cortical neural cultures were set up and maintained according to^30,76^. At DIV6, cultures were exposed to KP (50, 100, 400 nM) for 48h and then harvested for isolation of RNA. For immunostaining, the experiment was repeated in independent cultures using 400 nM KP. Cells were fixed with 4% paraformaldehyde and processed for immunolabeling for KCC2, neuronal synaptic marker, synaptophysin, and neuronal nuclear differentiation marker (NeuN), and then inspected and image-captured on a fluorescent confocal microscope (Zeiss LSM700).

### δ-catenin DNA constructs, transgenesis vectors

Human δ2-catenin isoforms (δ-cat, WT, and S276A point mutation) were cloned into pAAV-hSyn, tdTomato was used as control. AAV9 particles were packaged (Duke Viral Vector Core Lab). For i.t injections, a dilution of 10^12^ viral genomes/mL was used.

### *Kcc2* promoter luciferase reporter assays

After identifying TCF and Kaiso binding sites within the *Kcc2b* regulatory region (position −2052 bp to +476 bp relative to transcriptional start site (TSS)), these specific sites were deleted by PCR-based cloning. Following previous methods^18^, the WT promoter and the respective engineered mutant promoters were cloned into pGL4.17 to drive LUC. N2a cells were transfected with these constructs, co-transfected with Renilla LUC for normalization, and assayed for dual-LUC activity 24h after transfection. Mutant promoters were compared to WT, and the response to KP (400 nM) by the respective promoter was measured.

### Chemicals

Kenpaullone compound was synthesized by the Duke Small Molecule Synthesis Facility to >98% purity, verified by LC/MS. CHIR99201, and VU0240551 were obtained from Tocris. GW801372X, GW778894X, GW300660X, GW779439X, and GW305178X were supplied by the Structural Genomics Consortium (SGC) at UNC-Chapel Hill.

### Animals

C57bl/6J male mice (10-12 weeks old) were obtained from The Jackson Lab (Bar Harbor, ME). *Kcc2*-LUC mice were generated by the Liedtke Lab at Duke University and continued as a line within our mouse colony. All animal procedures were approved by The Duke University IACUC and carried out in accordance with the NIH’s Guide for the Care and Use of Laboratory Animals.

### RT-qPCR from cultured neuronal cells and microdissected spinal cord dorsal horn

oligodT-initiated reverse transcription of 1 μg total RNA, DNA-se treated, was subjected to RT-qPCR using primers specific for *Kcc2* for rat and *KCC2* for human sequences, normalized for neuronal β_III_-tubulin, as previously described^18^. For cultured neurons, total RNA was extracted from pelleted cells, and for spinal cord tissue it was extracted from microdissected lumbar spinal cord dorsal horn.

### Behavioral assessments

For pain-related assays, hind paw withdrawal in response to mechanical stimuli was assessed with von Frey hairs (vFH) using an automated vFH apparatus^77,78^ (Ji-Lab). Sensitization was implemented by sciatic nerve injury via ligation, using the PSNL and CCI well-established models^37,38^. For in vivo behavioral assays, mice were injected starting on day 1 after constriction injury with 10 or 30 mg/kg KP, intraperitoneally (i.p.), or intrathecally at 30 μg in 5 μL.

Intrathecal compound or viral vector injection was conducted as in^79^.

Vestibulomotor function, cerebellar coordination, and motor stamina were assessed using an automated rotarod (RR) (Ugo Basile, Italy)^41^. Animals were RR-trained pre-injection of KP, which was injected daily, and their stamina/ RR performance was recorded for two weeks.

Conditioned place preference (CPP) was conducted as in^80^ by recording choice behavior for one of two accessible chambers after 7 days of conditioning; animals were treated with KP at 30 mg/kg or vehicle in one chamber. CPP scores were calculated as post-conditioning time minus preconditioning time spent in the treatment-paired chamber.

### Bone cancer pain model

Murine lung carcinoma cell line LLC1 (ATCC CRL-1642) was injected into mouse femora following previous protocol^40^. Under brief general anesthesia, 2 × 10^5^ tumor cells were injected into the left femur’s cavity. Mechanical sensitivity was assessed as described above for nerve constriction injury. Osteolytic bone lesion was determined by faxitron, radiographs read by blinded investigators as described in^39,40^.

### Chromatin immunoprecipitation

ChIP assay was carried out as described previously^18^ using primary cortical neurons (0.7 × 10^6^).

### Cultured neuron immuno-cytochemistry

Immuno-cytochemistry labeling of cultured neuronal cells was carried out as previously described^18^ using antibodies specific for δ-cat, ß-cat, ß-tubulin_III_, and FLAG epitope. Imaging and morphometry of acquired images was conducted using a Zeiss LSM710 microscopy platform with Zen software.

### Spinal cord immuno-histochemistry

Immunohistochemical labeling of lumbar spinal cord was conducted as in^31,81,82^ using KCC2-specific antibodies.

### Chloride imaging

Chloride imaging of primary cultured cortical neurons was conducted as in^18,31,32^. Clomeleon ratiometric fluorescent chloride indicator protein was transfected into neurons and ratios were acquired on an Olympus BX10 microscope using RATIOTOOL software. Calibration experiments were conducted as in^18,32^.

### Spinal cord dorsal horn electrophysiology

Spinal cord slices (300-400 μm thickness) were prepared from male mice 5-7 weeks of age as in^83,84^. The gramicidin perforated patch method was applied to measure reversal potential for GABA of patched layer-II neurons; 1 mM GABA was applied with a puff-pipette^85,86^.

### Drug affinity responsive target stability (DARTS) assay

DARTS assay was conducted following^44^ using 20μM KP as “bait” and LC-MS/LC to identify proteins bound to KP. The assay was conducted in both the presence and absence of detergents NP40 and N-dodecyl-b-D-maltoside.

### Kinome analysis

Cultured rat primary cortical neurons were treated with either vehicle (DMSO 0.1%) or 1 μM KP for 1h/24h. Protein extract was purified, trypsin-digested, and enriched for phosphopeptide via TiO2 resin. Identity and quantity of enriched phosphopeptides were determined using analytical LC-MS/MS, and then samples were analyzed for differential expression (KP vs. vehicle treatment) for both time points.

### Statistics

All data are expressed as mean ± SEM. Differences between groups were evaluated using two-tailed Student’s *t* test (experimental against sham control), or in the case of multiple groups, one-way ANOVA followed by post-hoc Tukey test. For time-courses, a two-way ANOVA followed by post-hoc Bonferroni test was used. The criterion for statistical significance was p < 0.05.

## Supplementary Materials

**Suppl. File S1: Excel sheet #1. Compound screening results**

**Suppl. File S2: Excel sheet #2. DARTS assay**

**Suppl. File S3: Excel sheet #3. Phosphoproteomics data**

**Detailed Methods**

**Supplementary Tables**

**Supplementary Figures**

## Detailed Methods

### Screening in primary cortical neurons from *Kcc2*-LUC transgenic mice

Transgenic mice that express red-shifted luciferase (LUC) under the control of the *Kcc2* promoter (−2052/+476, described in^18^), inserted into the Rosa26 locus, were previously described by our laboratory^30,31^. We generated primary cortical neuronal cultures from newborn (p0) mice of this line following^30,31^.

Cytosine arabinoside (2.5 μm) was added to cultures on the second day after seeding [2 d *in vitro* (DIV)] to inhibit the proliferation of non-neuronal cells. Cell suspension was plated at a density of 1 × 10^6^ cells/ml onto 24-well tissue-culture dishes coated with poly-d-lysine. Cortical neuronal cultures prepared by this method yielded a majority population of neuronal cells, with negligible glia contamination, as evidenced by the absence of GFAP by Western blotting.

After a week in culture, neurons were treated with compounds (100 nM for 48h). LUC activity was then determined for each compound. Culture supernatant was removed and cells were lysed with 150 μl lysis buffer (Targeting Systems CA, USA cat. CLR1). LUC activity was measured with a *Red*Luciferase Assay kit (Targeting Systems cat. FLAR) according to the manufacturer’s instructions. A Veritas microplate luminometer was used to measure luminescence; for each treatment triplicates of 40 μl cell lysates (from each 24-well tissue-culture dish) were used and 25μl substrate was injected per well. To evaluate the quality of screening methodology, Z’ factor^87^ was ascertained using 0.5% (v/v) DMSO as negative control and Trichostatin A as positive control. Relative Light units (RLU) for each treatment was derived from Light Units (compound treatment)/ vehicle treatment (0.5% DMSO).

The primary screen encompassed three levels of LUC measurements. The first level of screening yielded a total of 137 compounds with cut-off RLU >125% LUC activity. These 137 compounds were subject to second round of screening, which was carried out in duplicate independent assays to yield the top 103 compounds sorted for highest RLU. The 103 compounds were then subjected to a third round of screening carried out in duplicate independent assays to yield the top 40 compounds which were again ranked. The best 22 of these 40 compounds (ranked for highest RLU activity) were subjected to secondary screening: Cultured neurons were treated with the 22 compounds; RT-qPCR and Clomeleon imaging methodologies^18^ were used to determine effects of compounds on *Kcc2* mRNA expression and [Cl-]i, respectively. Each compound was ranked based on composite scores from primary and secondary screening.

### Human neuronal cultures

The neuron-enriched cultures were established from fetal cortical specimens at 15–20 weeks of gestation. The protocols for tissue processing complied with all federal and institutional guidelines. The cultures were plated on PEI (polyethyleneimine solution) substrate and maintained in Neurobasal media supplemented with B27 as described^30,76^.

For RT-qPCR, the cell pellets were collected after treatments with vehicle (0.1% DMSO) or KP (50, 100, and 400 nM) for 48h starting at day 6 in vitro.

For immunostaining, the cultures were treated for 2 – 4 days with vehicle or 400 nM KP, fixed with 4% PFA at day 10, and processed for double-staining with anti-KCC2 and anti-synaptophysin or anti-NeuN (see antibody table). The 0.34 micrometer confocal slices through entire cell layers at different optical fields (n = 17 – 28 for each group) of the fixated cultures were acquired using a Zeiss LSM700 confocal microscope.

Image Z-stacks were analyzed using Imaris 9.2.1 software. Optical density intensity sum values for each stack/channel were divided by corresponding data volumes and the resulting values were normalized to the number of specifically labeled cells within each stack. There were no significant differences in the number of cells between vehicle and KP-treated groups.

### δ-catenin DNA constructs, transgenesis vectors

Plasmid containing human δ-catenin (CTNND2, NM_001288717) open reading frame was obtained from GeneCopeia (EX-A4285-M02) and cloned into pCMV-ENTER vector (Origene PS100001). Site-directed mutagenesis using Phusion DNA Polymerase enzyme (Thermofisher F549L) in conjunction with complementary primers bearing the specific mutation were used to generate the S276A δ-catenin mutation S276A. PCR was followed by Dpn1 enzyme digestion to remove parental plasmid DNA. All constructs were verified by sequencing. pCS-CMV-tdTomato plasmid was obtained from Addgene (cat. #30530).

Plasmid pAAV-hSyn-eNpHR 3.0-EYFP from Addgene (26972) was cut with Age I and Hind III enzymes to excise the eNpHR 3.0-EYFP open reading frames. The control tdTomato open reading frame as well as the wild-type and mutant delta catenin open reading frames were generated with Age I and Hind III ends by PCR and subsequently inserted into the Age I / Hind III digested pAAV-hSyn plasmid. Orientation and sequence fidelity in the final constructs were verified by PCR and sequencing. AAV9 particles were packaged by the Duke University Viral Vector Core facility and were used at a titer of 10^12^ viral genome copies per mL.

### *Kcc2* promoter luciferase reporter assays

A fragment of the mouse *Kcc2* gene promoter (position −2052kbp to +476kbp) was amplified from genomic DNA prepared from cultured mouse primary glial cells. A 2.5kb PCR fragment was cloned into the pGL4.17-Basic Vector (Promega) to generate the *Kcc2* promoter reporter construct. TCF and Kaiso binding sites were identified in this fragment. Using wild-type construct pGL4.17-*Kcc2* as a template, site-directed mutagenesis using Phusion DNA Polymerase enzyme (Thermo Fisher F549L) in conjunction with complementary primers bearing the specific mutation were used to mutate the Kaiso and TCF DNA-binding sites. PCR was followed by Dpn1 enzyme digestion to remove parental plasmid DNA. All constructs were verified by sequencing.

N2a cells were grown to 90% confluency in 24-well dishes in 0.4 mL of medium (DMEM, 2% Fetal Bovine Serum, 2 mM glutamine, 1% Non-essential amino acids, and 1% Penicillin/Streptomycin). Cells were transiently transfected using TurboFect reagent (Thermo Fisher R0531), with 500 ng of the pGL4.17-constructs plus 20 ng of the control Renilla plasmid (Promega, E2231) to normalize for transfection efficiency. Twenty-four hours after transfection, luminescence was measured using the Dual-Luciferase^®^ Reporter Assay System (Promega) in a microplate luminometer (Veritas, Turner Biosystems). Mutant promoters were compared to WT, and the response of the respective promoter to KP (400 nM) was measured. Three independent transfection experiments were carried out and LUC assays were done in triplicates for each transfection. RLU is expressed as firefly luciferase activity relative to Renilla LUC activity.

### RT-qPCR

Total RNA was isolated from cultured cell samples using Directzol RNA miniprep kit (ZymoResearch). The protocol includes DNAse digestion to exclude genomic DNA from preparations. Total RNA (1μg)was reverse transcribed using oligo primers (dT) and SuperScriptIII first-strand synthesis kit (Invitrogen). Gene expression was assessed by quantitative real-time PCR using 2× SYBR Green Master Mix (Qiagen) and a three-step cycling protocol (anneal at 60°C /elongate at 72°C, denature at 95°C). Specificity of primers was verified by dissociation/melting curve for the amplicons when using SYBR Green as a detector. All reactions were performed in triplicates. The amount of target messenger RNA (mRNA) in the experimental group relative to that in the control was determined from the resulting fluorescence and threshold values (Ct) using the ΔΔCt method. β_III_-tubulin was used as housekeeping gene.

### Behavioral assessments

For pain-related behavior, mechanical allodynia was assessed with von Frey filaments (Ugo Basile, Italy). Animals were habituated to the testing environment daily for at least 2 days before baseline testing. The room temperature and humidity remained stable for all experiments. The mice were placed on a 5×5-mm wire-mesh grid floor in individual compartments to avoid visual stimulation and allowed to adapt for 0.5 h prior to the von Frey test. The von Frey filament was then applied to the middle of the plantar surface of the hind paw, perpendicularly, with a series of von Frey hairs with logarithmically increasing stiffness (0.02–2.56g, Stoelting). The withdrawal responses following the hind paw stimulation were measured at least three times. We determined the 50% paw withdrawal threshold by up-down method^78^.

For assessment in rotarod (RR), all animals received training prior to experiments; mice were placed on the RR apparatus set in an accelerating rotational speed mode (3–30 rpm, 300 s max) per trial. Following training, the average time to fall from the rotating cylinder over three trials was recorded as baseline latency (4-40rpm, 300s max/trial). Mice were injected daily with either vehicle or drug compounds before RR tests. Latency to fall was measured (4-40rpm, 300s max/trial (inter-trial interval is at least 15 min). The average latency to fall from the rod was recorded for each animal.

Conditioned place preference (CPP) was conducted using a CPP box, which consists of two conditioning chambers distinguished by visual and sensory cues, along with a small buffering chamber. All mice received a 3-day preconditioning habituation period with free access to both conditioning chambers and the time spent in each chamber was recorded for 15 min on day 3 after habituation. On conditioning days (day 4-10), mice first received the vehicle control (i.p. 5% DMSO, 5%Tween-80 in normal saline) paired with a randomly chosen chamber in the morning. After 4 hours, mice received KP (i.p. 30mg/kg) or vehicle, paired with the other chamber. During the conditioning, mice were allowed to stay only in the paired chamber for 15 min without access to other chambers. On test day (d11), mice were placed in the buffering chamber with free access to both conditioning chambers and choice behavior was recorded for 15 min. The CPP scores were calculated as post-conditioning time minus preconditioning time spent in the paired chamber.

### Bone cancer pain model

Murine lung carcinoma cell line LLC1 (ATCC CRL-1642) was digested with 0.25% trypsin and suspended at 5×10^7^/ml cells in PBS. Following previous protocol^40^, mice were anesthetized with 3% isoflurane (oxygen flow: 1.0 L/min). The left leg was shaved, and the skin was disinfected with 10% povidone-iodine and 75% ethanol. A 0.5-1cm superficial incision was made near the knee joint to expose the patellar ligament. Then a 25-gauge needle was inserted at the site of the intercondylar notch of the left femur into the femoral cavity and the needle was then replaced with a 10 μL microinjection syringe containing 4 μL suspension of tumor cells (2 × 10^5^) and 2 μL absorbable gelatin sponge solution for closure of the injection site. The contents of the syringe were slowly injected into the femoral cavity (2 min). To further prevent leakage of tumor cells from the bone’s cavity, the injection site was sealed with silicone adhesive at the periost level. Animals with surgery related movement dysfunction or with mis-targeted tumor cell injection were excluded.

### Radiography of bone imaging

Osteolytic bone destruction was assessed by Faxitron (Faxitron Bioptics, Tucson, Arizona). Radiographs of tumor-bearing femora were rated following a 0-5 score scale as previously described^39,40^ by blinded readers: 0, normal bone without signs of destruction; 1, one to three radiolucent lesions indicative of bone destruction; 2, increased number of lesions (three to six lesions) and loss of medullary bone; 3, loss of medullary bone and erosion of cortical bone; 4, full-thickness unicortical bone loss; 5, full-thickness bicortical bone loss and displaced skeletal fracture.

### Chromatin immunoprecipitation

ChIP assay was carried out as described previously^18,30^. Primary cortical neurons (0.7 × 10^6^) were used for each ChIP experiment. Cells were crosslinked with 1% formaldehyde for 30 min, washed twice with cold PBS, resuspended in lysis buffer [1%SDS, 10 mm EDTA, and 50 mm Tris-HCl, pH 8.0, with protease inhibitor cocktail (Roche)], and sonicated for 15 s pulses. The lysates were clarified by centrifugation at 10,000 rpm for 10 min at 4°C in a microcentrifuge. One-tenth of the total lysate was used as input control of genomic DNA. Supernatants were collected and diluted in buffer (1% Triton X-100, 2 mm EDTA, 150 mm NaCl, 20 mm Tris-HCl, pH 8.0, and protease inhibitor cocktail) followed by immunoclearing with 1 mg of salmon sperm DNA, 10 ml of rabbit IgG, and 20 ml of protein A/G-Sepharose (Santa Cruz Biotechnology) for 1h at 4°C. Immunoprecipitation was performed overnight at 4°C with 2 mg of each specific antibody. Precipitates were washed sequentially for 10 min each in TSE1 buffer (0.1% SDS, 1% Triton X-100, 2 mm EDTA, 150 mm NaCl, and 20 mm Tris-HCl, pH 8.0), TSE2 (TSE1 with 500 mm NaCl), and TSE3 (0.25 m LiCl, 1% NP-40, 1% deoxycholate, 1 mm EDTA, and 10 mm Tris-HCl, pH 8.0). Precipitates were then washed twice with 10 mm Tris/0.1 mm EDTA, pH 7.8 and extracted with 1% SDS containing 0.1 m NaHCO3. Eluates were pooled and heated at 65°C for 4 h to reverse formaldehyde crosslinking. DNA fragments were purified with Qiagen Qiaquick kit. For ChIP PCR, 1 μl of a 25 μl DNA extraction was used.

### Immuno-cytochemistry of cultured neurons

Immunocytochemistry labeling of cultured neuronal cells was carried out as previously described^l8,30,31^. Primary antibodies are shown in the antibody table. Anti-KCC2 primary antibodies were validated with developing rat primary cortical neurons; we observed an increase in staining pattern that tightly matched increase of *Kcc2* mRNA expression^18,30,31^. Secondary antibodies used were goat anti-mouse IgG Alexa Fluor 594 (Invitrogen A11032) and goat anti-rabbit IgG Alexa Fluor 594 (Invitrogen A11012). DAPI stain was obtained from Sigma Aldrich (D9542). Stained cells were observed using an inverted confocal microscope (Zeiss LSM780).

We obtained stacks of images recorded at 0.35 μm intervals through separate channels with a 63x oil-immersion lens (NA, 1.40, refraction index, 1.45). Zen software (Zeiss) was used to construct composite images from each optical series by combining the images recorded through the different channels, and the same software was used to obtain Z projection images (image resolution: 1024 × 1024 pixels; pixel size: 0.11 μm). ImageJ was used for morphometry.

### Spinal cord immuno-histochemistry

All mice were deeply anaesthetized with isoflurane and then transcardially perfused with ice-cold 4% paraformaldehyde in 0.1 M phosphate buffer, pH 7.4 (4% PFA). Dissected spinal cord samples were then post-fixed overnight in 4% PFA at 4 °C, cryoprotected in a 20% sucrose solution in PBS at 4 °C, frozen in Tissue-Tek OCT (Sakura), and stored at −80 °C until sectioning. Samples were sectioned at 20 μm using a cryostat (Microm HM 505N). The sections were blocked with 2% bovine serum albumin (BSA) in PBS with 0.3% Triton X-100 (Blocking solution) at room temperature for 1h. The sections were treated with primary antibody in blocking solution at 4°C overnight. The sections were washed three times followed by secondary antibody treatment at 4°C for 2 hours. Anti-KCC2 antibody was validated as described above for immunocytochemistry. The goat anti-rabbit IgG Alexa Fluor 488 was obtained from Invitrogen (A-11008). Morphometry was conducted using ImageJ with region-of-interest Rexed laminae I-II.

### Drug Affinity Responsive Target Stability (DARTS) assay

Cultured primary rat cortical neurons were treated with either vehicle DMSO (0.1%) or 20 μM KP for 30h. Cells were lysed in ice-cold lysis buffer (Tris.Cl pH8 50mM, NaCl 150mM, NP40 0.5%, N-dodecyl-b-D-maltoside 0.5%, Phosphatase Inhibitor (Pierce #88667) and Protease Inhibitor (Roche #11836153001)). Protein concentrations were determined by Bio-Rad DC Protein Assay kit using bovine albumin as standard. All steps were performed on ice. Samples were warmed to room temperature and digested with pronase (final concentration 1:500) for 30 min at 30 °C. Digestion was halted using 0.5M EDTA. Only proteins not bound to KP were digested. The protein mixture was dialyzed using dialysis cassettes (Thermo Fisher 66203, 2K MWCO) and analyzed by LC-MS/MS method to identify proteins that are bound to KP, the latter step carried out in the Duke Proteomics Core Laboratory.

### Kinome analysis

Cultured rat primary cortical neurons were treated with either vehicle DMSO (0.1%) or 1 μM KP for 1h/24h. Cells were lysed in non-detergent-containing buffer (Tris.Cl pH8 50mM, NaCl 150mM, 0.5% Phosphatase Inhibitor (Pierce #88667) and Protease Inhibitor (Roche #11836153001)). 500 μL was removed and solid urea was added to a final concentration of 8M. Samples were sonicated for further solubilization. After clearing of insoluble material by centrifugation, protein concentration was measured by Bradford assay. 250 μg of total protein was removed from each sample and solubilization buffer was added to normalize all samples to 0.93 μg/μL protein. Samples were then spiked with bovine alpha-casein to 30 fmol/μg of total protein. Samples were reduced with 10 mM DTT at 32 °C for 45 min and then alkylated with 20 mM iodoacetamide at room temperature for 30 min. Samples were trypsin digested at 1:25 (enzyme-to-protein) overnight at 32 °C. Following acidification with TFA to pH 2.5, samples were subjected to a C18 solid-phase extraction cleanup. Eluted peptides were split 80% for phosphopeptide analysis and 20% reserved for unbiased differential expression. The phosphopeptide fraction (200 μg) was then frozen and lyophilized prior to phosphopeptide enrichment.

TiO2 Enrichment: Samples were resuspended in 65 μL of 1M glycolic acid in 80% MeCN/1% TFA and were enriched on TiO2 resin using a 10 μL GL Sciences microliter TiO2 spin tips following an established protocol (http://www.genome.duke.edu/cores/proteomics/samplepreparation/documents/GL_SpinColumnProtocol_bmr_ejs_mt_061713.pdf). After elution and acidification, samples were lyophilized to dryness and resuspended in 100 μL of 0.15% TFA in water. After cleanup using a C18 STAGE tip, and resuspension in 2% acetonitrile, 0.1% TFA, 10 mM citric acid samples were quantified.

Quantitative analysis of Phosphopeptide Enriched Samples: Quantitative LC-MS/MS was performed in singlicate (4uL=33% of the total sample each injection) for phosphopeptide-enriched samples using a nanoAcquity UPLC system (Waters Corp) coupled to a Thermo QExactive Plus high resolution accurate mass tandem mass spectrometer (Thermo) via a nanoelectrospray ionization source. Briefly, the sample was first trapped on a Symmetry C18 300 mm Å~ 180 mm trapping column for 6 min at 5l/min (99.9/0.1 v/v water/acetonitrile 0.1% formic acid), after which the analytical separation was performed on a 1.7 μm Acquity BEH130 C18 75 mm Å~250 mm column (Waters Corp). Peptides were held at 3% acetonitrile with 0.1% formic acid for 5 min and then subjected to a linear gradient from 3 to 30% acetonitrile with 0.1% formic acid over 90 min at a flow rate of 400 nL/min at 55°C. Data collection on the QExactivePlus mass-spec was performed in a data-dependent acquisition (DDA) mode following protocol of the manufacturer.

### Chloride imaging

We followed methodology described previously^18,31,32^. A Clomeleon expression plasmid was transfected into primary cortical neurons by electroporation (Amaxa Nucleofector Device). Transfected neurons were verified by yellow fluorescent protein (YFP) fluorescence, and ratiometric images (excitation at λ = 434 nm, dual emission at λ = 485 and 535 nm; for resting chloride, six stable frames at a rate 12 of per minute were captured, which were averaged) were acquired using RATIOTOOL program. Calibration of Clomeleon signals (535 nm/485 nm emission ratio) was performed by using tributyltin-nigericin to establish a standard curve (Pond et al., 2006), which was then normalized for measured intraneuronal pH to take into account the pH sensitivity of Clomeleon.

### Spinal cord dorsal horn electrophysiology

For spinal cord slice preparation, adult (5-7 weeks) male mice were anesthetized with urethane (1.5-2.0 g/kg, i.p.). The lumbosacral spinal cord was microsurgically removed and submerged into ice-cold dissection media which was saturated with 95% O_2_ and 5% CO_2_ at room temperature. After extraction and still under anesthesia, animals were euthanized. Transverse slices (300-400 μm) were cut using a vibrating microslicer (VT1200s Leica). The slices were incubated at 32°C for at least 30 min in regular artificial cerebrospinal fluid (aCSF), equilibrated with 95% O_2_ and 5% CO_2_.

The following solutions were used: Dissection solution: Sucrose 240 mM, NaHCO_3_ 25 mM, KCl 2.5 mM, NaH_2_PO_4_ 1.25 mM, CaCl_2_ 0.5 mM, MgCl_2_ 3.5 mM^88^. Regular artificial cerebrospinal fluid (ACSF): NaCl 117 mM, KCl 3.6 mM, MgCl_2_ 1.2 mM, CaCl_2_ 2.5 mM, NaHCO_3_ 25 mM, NaH_2_PO_4_ 1.2 mM, glucose 11 mM. The pH value of ACSF or dissection solution was adjusted to 7.4 when saturated with the gas. Normal intrapipette solution (pH 7.2 and 310 mOsm): K-methylsulfate 115 mM, KCl 25 mM, MgCl_2_ 2 mM, HEPES 10 mM, GTP-Na 0.4 mM and Mg-ATP 5 mM.

Electrophysiological recordings were conducted as follows. A slice was placed in the recording chamber and completely submerged and superfused at a rate of 2-4 ml/min with aCSF saturated with 95% O_2_ and 5% CO_2_ at room temperature. Perforated patch-clamp was used to avoid alteration of the [Cl-]i. To measure the chloride equilibrium potential (*E*_Cl_), gramicidin D (80 μg/mL with 0.8% DMSO final concentration, from an 8 mg/mL stock in DMSO) was added to the intrapipette solution, and 6-cyano-7-nitroquinoxaline-2,3-dione (CNQX, 10 μM), D_L_-2-amino-5-phosphonovaleric acid (APV, 50 μM), and tetrodotoxin (TTX, 0.5 μM) were added to the aCSF solution. The tip of the patch pipette was filled with the normal intrapipette solution while the rest of the pipette contained the gramicidin-containing solution. After forming a seal on the membrane, we waited ~30 min for the gramicidin to induce sufficient cation-selective pores in the membrane and lowered the series resistance to below 100 MΩ. Membrane potential measurements were corrected for liquid junction potential, which was measured as in^86^. GABA (1 mM) was puffed locally and instantaneously, and the puff pipette was aimed toward the recording pipette. To determine the reversal potential of GABA-evoked currents, voltage ramps were applied from +8 to −92 mV over 200 ms at a holding potential of −42 mV. Since the voltage ramp might elicit a basal current, a control voltage ramp was applied, and 1 min later GABA was puffed followed by another voltage ramp^85^. The reversal potential was analyzed as in^85^.

Signals were acquired using an Axopatch 700B amplifier and analyzed with pCLAMP 10.3 software. Only neurons with resting membrane potential < −50 mV and stable access resistance were included.

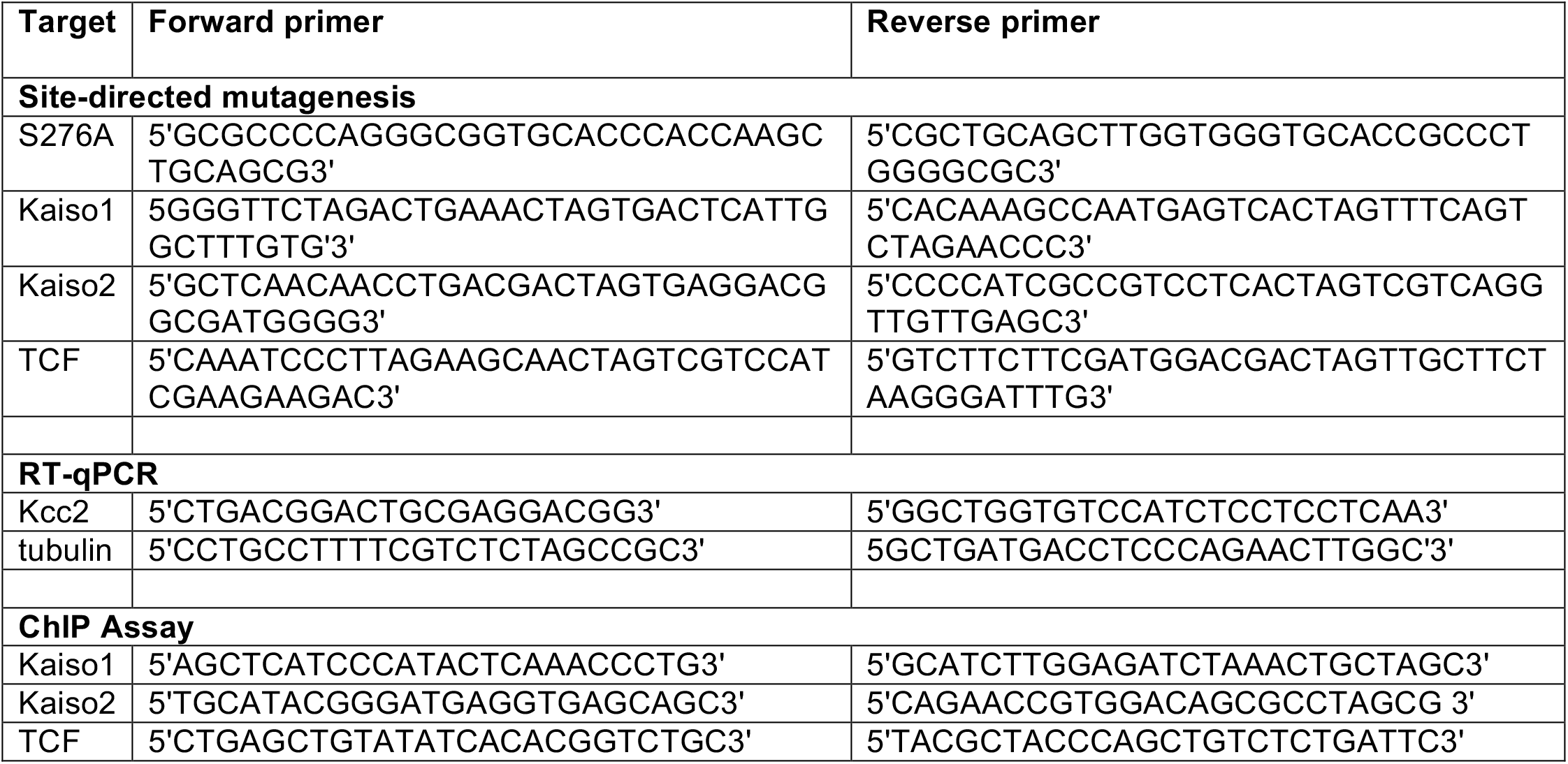
Supplementary Table of PCR Primers.

Supplementary Table of primary antibodies.

**Supplementary Table of primary antibodies**
| Antibody | Source |
| --- | --- |
| anti-KCC2 | Millipore (07-432) (mouse spinal cord)<br>ThermoFisher Scientific (PA5-78544) (human, mouse, rat ICC) |
| anti-$\beta_{III}$ Tubulin | Abcam (mouse, ab78078; rabbit, ab229590) |
| anti-β-catenin | SigmaAldrich (PLA0230) |
| anti-δ-catenin | SigmaAldrich (MABN2254) |
| anti-Synaptophysin | ThermoFisher (MA1-213) |
| anti-FLAG | SigmaAldrich (F3165) |
| anti-NeuN | BioLegend (834501) |

## Supplementary Figures

**Fig. S2.**
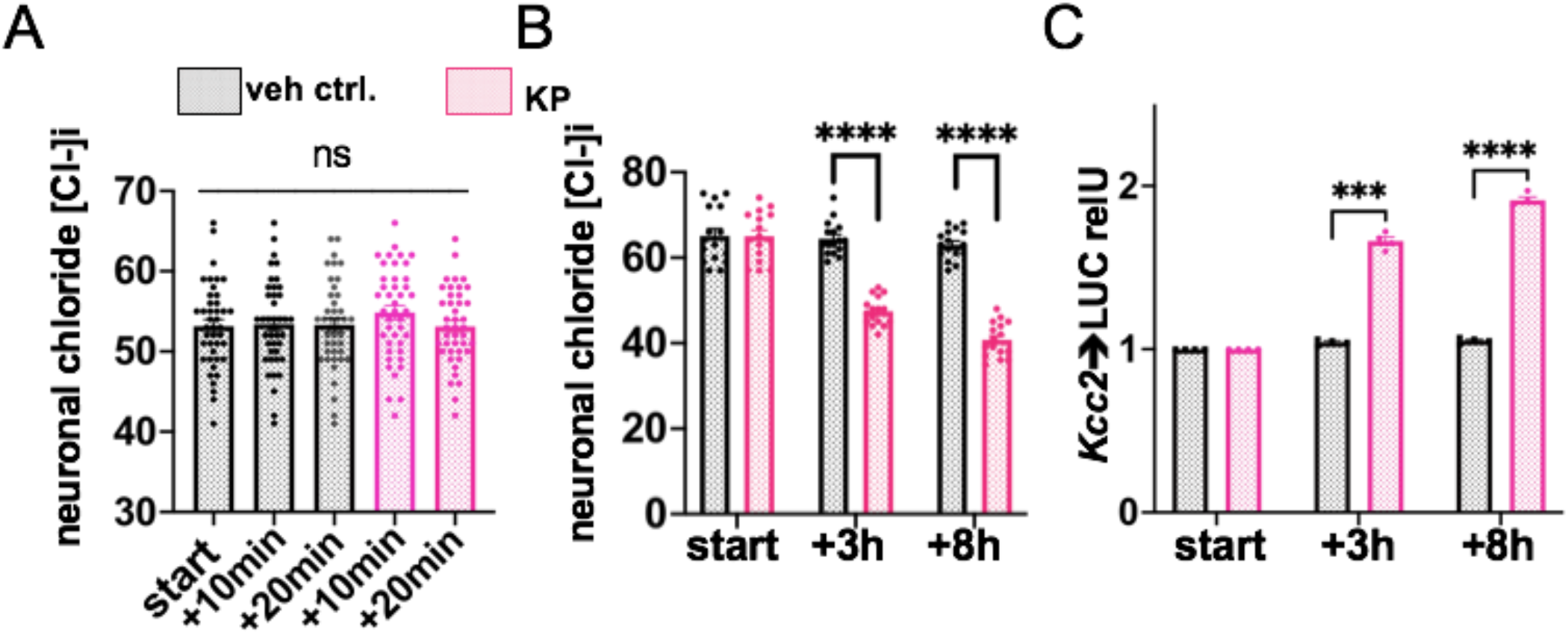
Kenpaullone does not enhance KCC2 chloride transporter mediated chloride efflux, affects Kcc2 expression and function within 3h. **A)** Rat primary cortical neurons, n=42 neurons/group. No difference was observed in neuronal chloride levels, measured with clomeleon fluorescent indicator (main ms., Fig. 2C) after KP treatment vs vehicle at the 10 min and 20 min time-points. **B)** Rat primary cortical neurons, clomeleon-based measurement of [Cl-]i, n=16 neurons/group. Note significant decrease of [Cl-]i at the 3h time-point, more pronounced at 8h. ****p<0.001, 2-way ANOVA. **C)** Rat primary cortical neurons, transfected with renilla luciferase, driven by *Kcc2* promoter, as in^18^. n=4 independent cultures. Note significantly increased activation of the *Kcc2* promoter at 3h, more pronounced at 8h. ***p<0.001, ****p<0.0001, 2-way ANOVA.

**Fig. S3.**
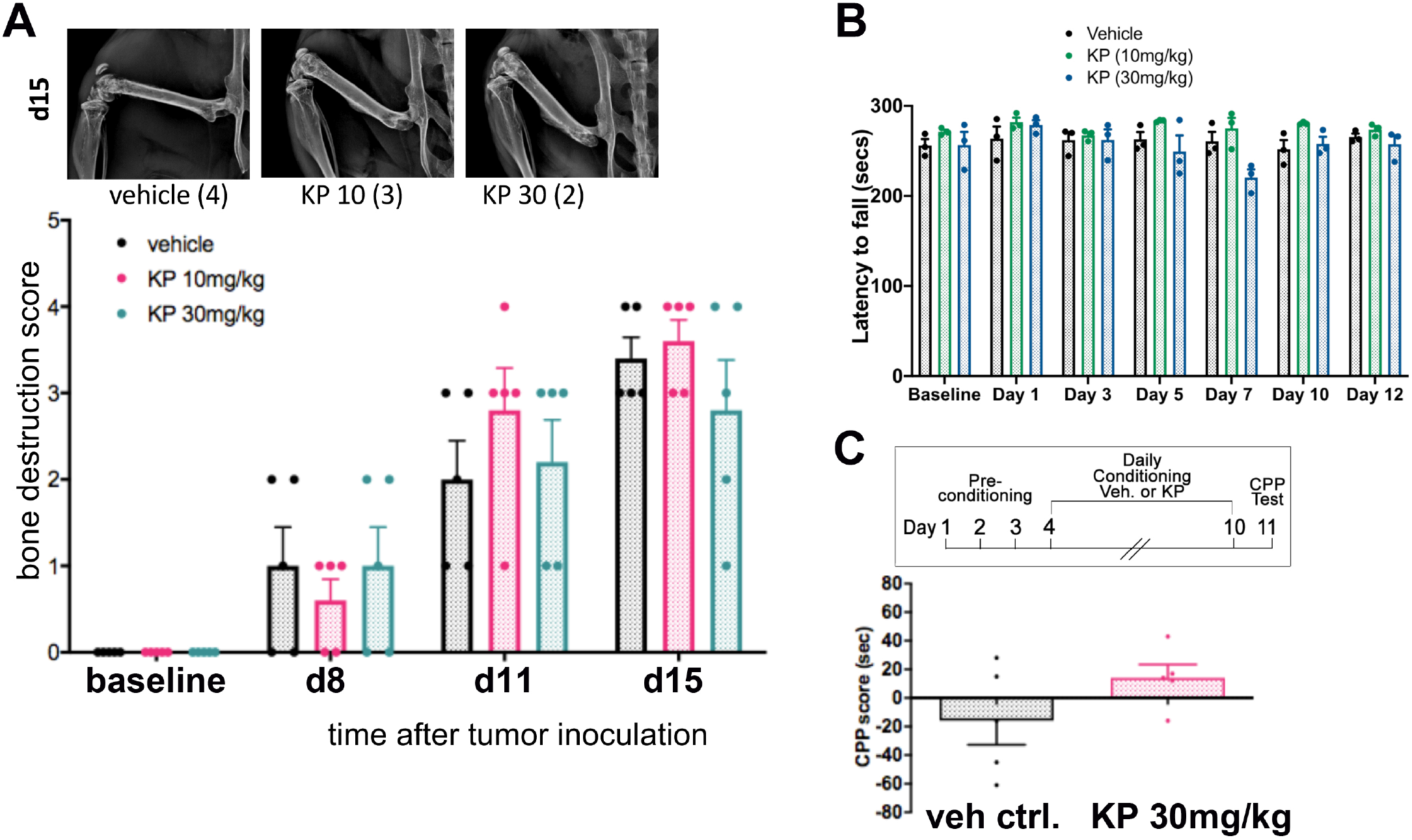
Kenpaullone is not affecting lytic bone lesions in a bone cancer model, is not affecting Rotarod-performance and conditioned place preference. **A)** Kenpaullone did not contain growth of mouse LLC lung tumor cells infused into the femur Osteolytic damage to femur (exemplars upper panel, at d15, for each group, with assessed severity) was not contained at any time point, not at 10 nor at 30 mg/kg bw KP, n=5 mice per group. **B)** Effect of KP on motor function and coordination of mice – rotarod assay. Bar diagram showing the mean of elapsed time on the rotarod. KP did not affect motor stamina and coordination in mice, with the one-time exception of high-dose KP on day 7. n=3-4 mice per group. **C)** KP does not evoke conditioned place preference (CPP). Timeline for CPP assay shown on top. Bottom: No significant difference in CPP scores between KP and vehicle; n=5 mice/group.

**Fig. S4.**
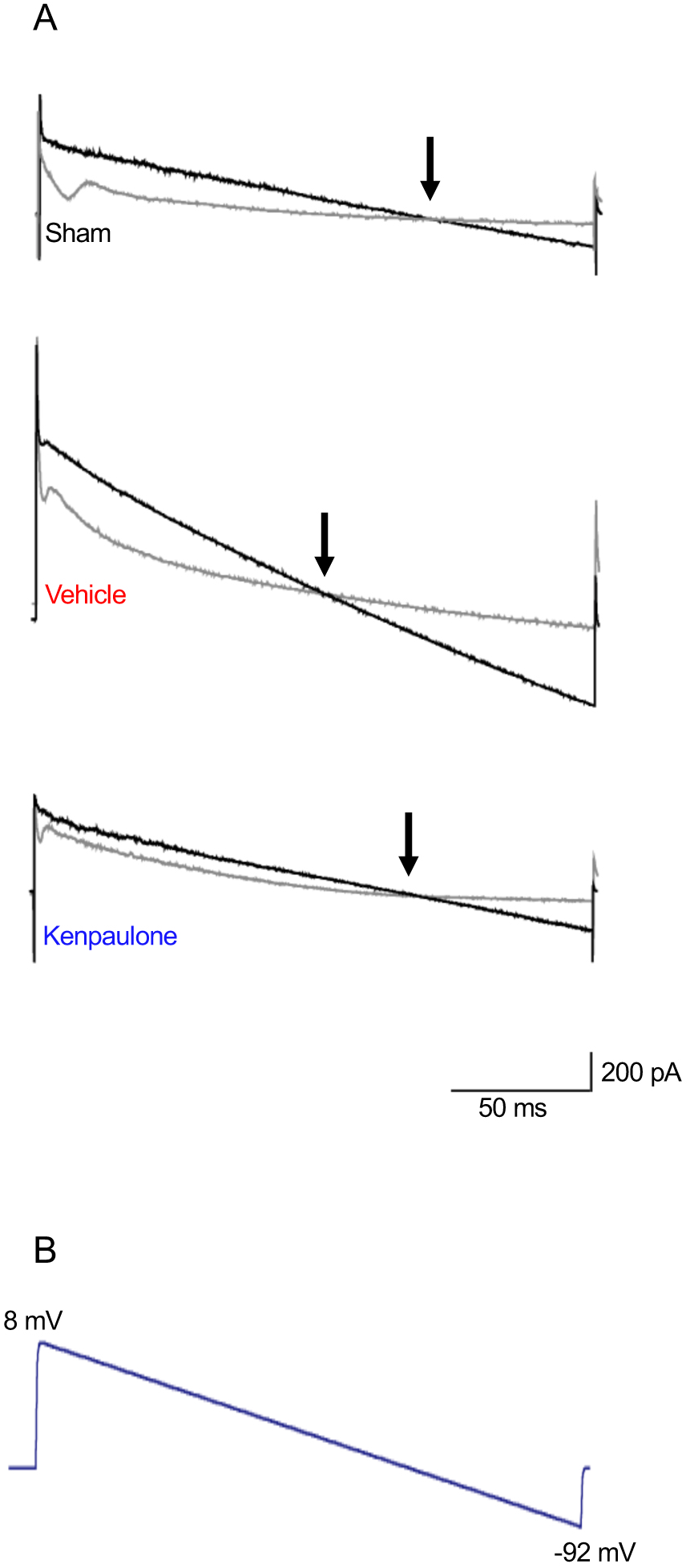
Electrophysiological recordings from spinal cord dorsal horn. **A)** Current responses of layer-II neurons to a voltage ramp from +8 to −92 mV (shown in panel **B**) in control (grey traces, obtained before GABA puff), or at the end of a puff of GABA (black trace) in sham, PSNL+vehicle and PSNL+KP groups. Reversal potential of the GABA-evoked current is at the voltage where the grey and black traces intersect (arrow).

**Fig. S6.**
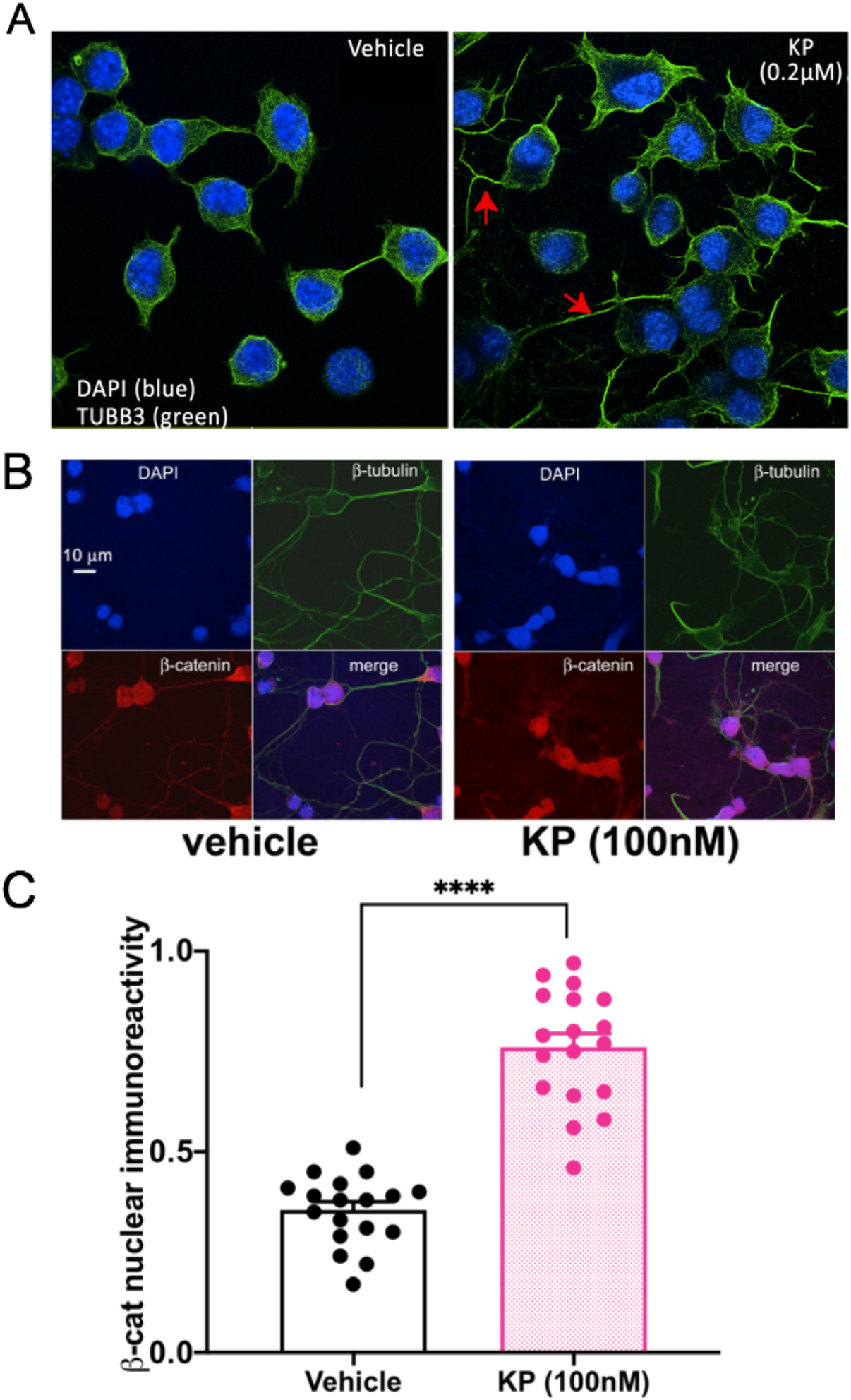
Kenpaullone enhances neuronal differentiation and facilitates β-catenin translocation to the nucleus. **A)** neuronalization of N2a cultured cells in response to KP (0.2μM for 3 days), note increased process formation and expression of neuronal β_III_-tubulin (green; red arrows). **B)** micrographs: Representative immuno-labeling of neuronal β_III_-tubulin (green) and β-catenin (red) before (left-hand panels) and after KP treatment (right-hand panels) in primary cortical neurons from rat. DAPI stain is used to define nuclear compartment for morphometric assessment. **C)** KP significantly increases β-catenin translocation into the nucleus; n=18 neurons/group.

**Fig. S7.**
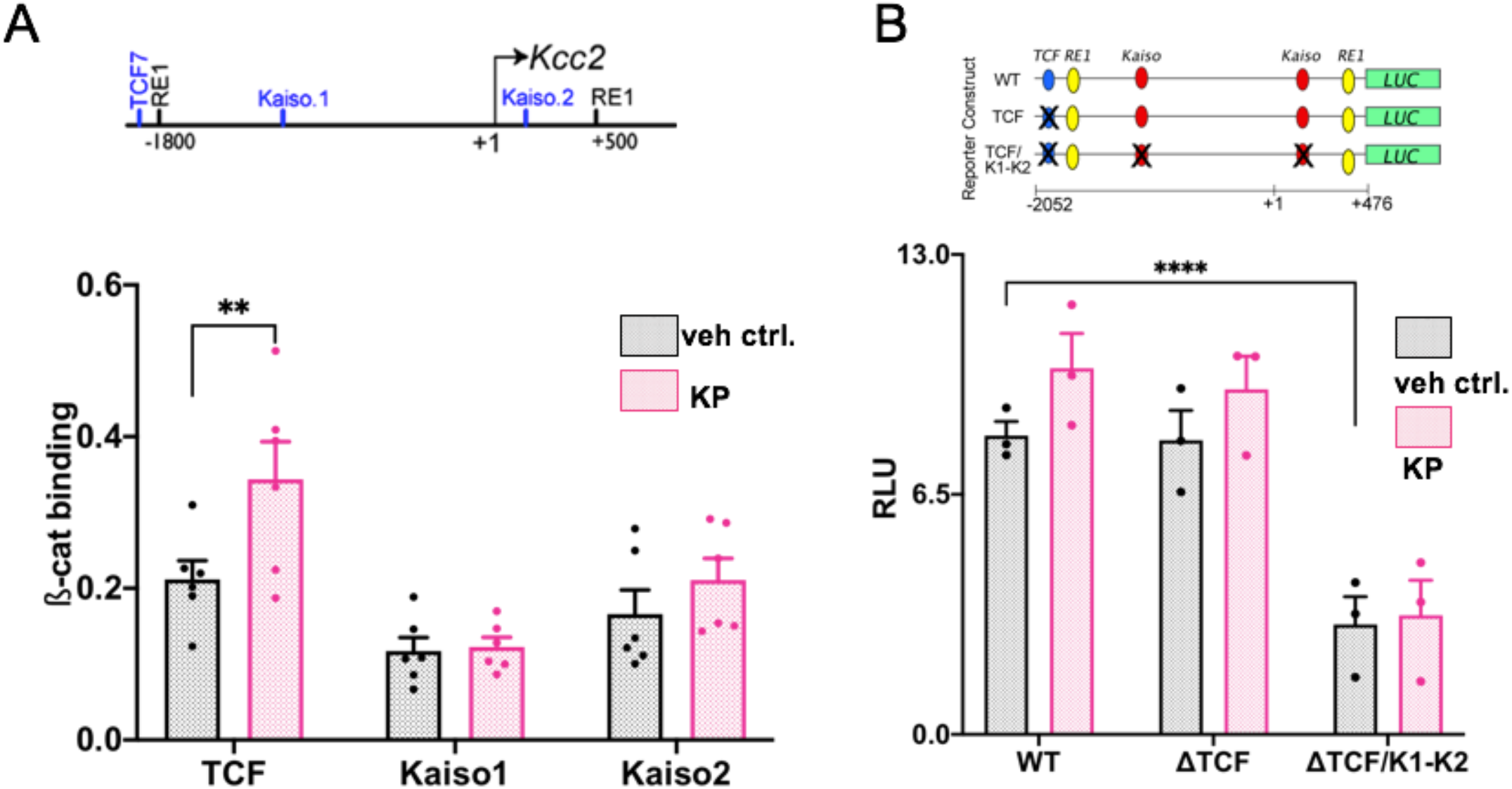
Kenpaullone increases ß-catenin binding to *Kcc2* promoter, relevance TCF site. **A)** KP increases ß-cat binding to the TCF site in the *Kcc2* promoter. Upper panel: Structure of mouse-*Kcc2* gene encompassing 2.5kb surrounding the TSS (+1), as in Fig. 7A. Bottom, bar diagram: Chromatin immuno-precipitation using anti-ß-cat antibody in primary rat cortical neurons reveals binding of ß-cat to TCF, to minor degree to Kaiso1, 2. KP treatment significantly increases binding of β-cat to the *Kcc2* promoter on the TCF binding site. n=6 independent rat primary neuron cultures were subjected to ChIP, **p<0.01 t-test, KP-treated vs vehicle. **B)** Relevance of TCF DNA binding site on regulation of *Kcc2*→LUC N2a differentiated cells were used as shown in Suppl Fig. 6A. Top panel: mouse *Kcc2* promoter constructs, as in Fig. 7B. Bottom, bar diagrams: KP did not increase promoter activity for any of the constructs. Triple-deletion of TCF and both Kaiso sites rendered the *Kcc2* promoter very low in activity, compared with WT (without KP), at <1/3 of its activity, which is less than the ΔΔKaiso1,2 deletion construct (Fig. 7B; <1/2). n=3 independent cultures, **** p<0.0001 WT vs triple-deletion construct, 1-way ANOVA.

